# Characterization of an immunodeficiency-associated EZH2 variant

**DOI:** 10.64898/2025.12.08.692979

**Authors:** Francisco Perez de los Santos, Lily Beck, Emily DeCurtis, April M. Griffin, Maggie M. Balas, Charlotte Marchioni, Attila Kumanovics, Matthew R. G. Taylor, Kimberly R. Jordan, Charles H. Kirkpatrick, Aaron M. Johnson

## Abstract

Regulation of gene expression is central to the development of immune cells and their ability to respond to infection. As part of a clinical evaluation, we identified two sisters with recurrent infections, hypogammaglobulinemia, and memory B cell deficiency, diagnosed as common variable immunodeficiency. Whole exome sequencing identified a heterozygous variant (Leu50Ser, L50S) in a conserved region of EZH2, the catalytic subunit of the epigenetic gene repressor Polycomb Repressive Complex 2 (PRC2). EZH2-catalyzed histone H3 lysine 27 methylation (H3K27me) in bulk was not overall significantly disrupted by this variant, in patient samples or cell lines expressing EZH2-L50S. EZH2-L50S protein is expressed similar to wild-type and can form PRC2. However, we find that specific genomic regions that normally have high wild-type levels of H3K27me3 are deficient in the L50S context, particularly around gene promoters. EZH2-L50S is still recruited to these sites, but is not as active. Using recombinant purified PRC2, we determine that L50S affects methylation of nucleosomes and disrupts allosteric stimulation that normally amplifies H3K27me3, consistent with the location of L50 in the allosteric regulatory region of PRC2. Thus, variation of EZH2 L50, occurring at low frequency in the population may interfere with normal B cell gene expression patterns, contributing to immunodeficiency. This study has implications for genetic variation in PRC2 in the general population.

## Introduction

Epigenetic regulation of gene expression by chromatin-based mechanisms is essential to development and cell differentiation in response to stimuli in adult tissues of the body. One major pathway for chromatin regulation is through the Polycomb group proteins that induce the formation of facultative heterochromatin^1^. Polycomb proteins exist in two primary complexes, polycomb repressive complex 1 and 2 (PRC1 and PRC2). Both complexes have histone modification activity, both can be recruited to chromatin by separate mechanisms to trigger facultative heterochromatic gene repression, and each can promote the other’s activity. PRC2 is solely responsible for catalyzing histone H3 lysine 27 (H3K27) methylation via the histone methyltransferase subunit EZH2, or its paralog EZH1 in some contexts. H3K27 methylation triggers heterochromatin formation as a preferred binding site for multiple “reader” subunits of PRC1, which can support chromatin compaction and gene repression. PRC2 is essential in early development, both to initially silence cell type-specific genes and also to turn off stem cell genes during differentiation. Later, in adult tissues PRC2 functions in two capacities: 1) maintaining previously-established polycomb repressed regions of the genome. This activity is carried out primarily by EZH1-containing PRC2, which is specialized for this function, rather than establishing new repression; 2) In a subset of tissues that go through major differentiation and proliferation programs, EZH2 is induced and EZH2-containing PRC2 is an important regulator. For example, both B cell differentiation and function require EZH2^2–5^. Loss of EZH2 leads to impaired development of B cells^2^. Activated mature B cells also express high levels of EZH2 and loss of EZH2 leads to functionally deficient memory B cells, including a decrease in germinal centers and antibody secreting cells and failing to generate a robust recall response^4–6^. On the other hand, increased EZH2 function in B cells due to somatic gain-of-function mutations leads to B cell lymphomas^7^.

The histone modification that PRC2 catalyzes, H3K27 methylation, not only promotes PRC1 activity, but it also is an allosteric stimulator of PRC2’s own activity^8^. This allosteric activation occurs through the PRC2 subunit EED, which, together with EZH2 and SUZ12, form the core subunits of PRC2^9^. EED can bind to tri-methylated H3K27 and this binding is transmitted to the active site of EZH2 to increase methylation of additional H3K27^10^. This mechanism is essential for all PRC2 activity, since mutation of the EED pocket that binds methylated H3K27 and disruption of the allosteric network result in severe loss of H3K27 methylation^8,10,11^. The allostery between EZH2 and EED is disrupted in the developmental skeletal overgrowth of Weaver Syndrome, where mutations in both genes are found^12–14^.

Here, we present a study that began with treatment of a pair of siblings who developed antibody deficiency and B cell loss during adult years. Whole exome sequencing disclosed that both have a heterozygous variant in EZH2, c.149T>C, present at 1 in ∼10-20,000 individuals, which will result in a substitution of leucine 50 to serine. In the remaining PBMC population, which is depleted of B cells, bulk H3K27 methylation is not significantly different from unaffected samples. Introduction of the L50S mutation into EZH2 in cells demonstrated stable L50S protein which was able to form PRC2. EZH2-L50S transgene expression led to a minor defect in bulk H3K27 tri-methylation levels. Genome-wide determination of H3K27me3 levels by CUT&Tag revealed that regions with normally high wild-type H3K27me3 levels were depleted in the EZH2-L50S context, though EZH2-L50S was still recruited to these regions similar to wild-type. Using purified PRC2 in biochemical assays, we found a decrease in nucleosome methylation and allosteric stimulation with EZH2-L50S compared to wild-type. Overall, our study shows that a genetic variant of EZH2 associated with reduced B cell function in adulthood has a hypomorphic molecular phenotype.

## Results

### Association of the L50S variant of EZH2 with a B cell immunodeficiency

In the course of the clinical practice of the late Dr. Kirkpatrick, a 38-year-old woman (I-2 in Figure 1A) was referred for evaluation of repeated respiratory infections including pneumonia. Later, her sister (I-4) was referred for evaluation of chronic hives and, over ∼5 years of follow up, began to have recurrent infections, including infectious sinusitis and bronchitis. The sisters were evaluated to have hypogammaglobulinemia, a severe defect in antibody production (Figure 1B) and were given the diagnosis of “common variable immune deficiency” (CVID). Both sisters were treated with immune globulin replacement therapy which has been beneficial. Prior to treatment, response after immunization was measured for three vaccines. Responses to *tetanus* toxoid and *H. influenza* b conjugate vaccines, were both well below the expected >=4-fold increase in titer in sera collected >=four weeks after immunizations. The pneumococcal vaccine (Pneumovax-23) response was also below normal, with 0 out of 14 (I-2) and 2 out of 14 (I-4) serotypes eliciting a >=4-fold response, whereas an expected normal response is at least 11 out of 14 (Figure 1C). Mononuclear cells were separated from heparinized whole blood by density gradient centrifugation and stained for lymphocyte type for general profiling (Figure 1D). All values for T-cells were in the normal range. The most striking abnormality observed was very low numbers of CD27+ memory B cells. These findings suggested that maturation of the peripheral B cell population had not progressed beyond the “naïve” (IgM+/IgD+CD27-) stage or there was a depletion of memory B cells.

**Figure 1.**
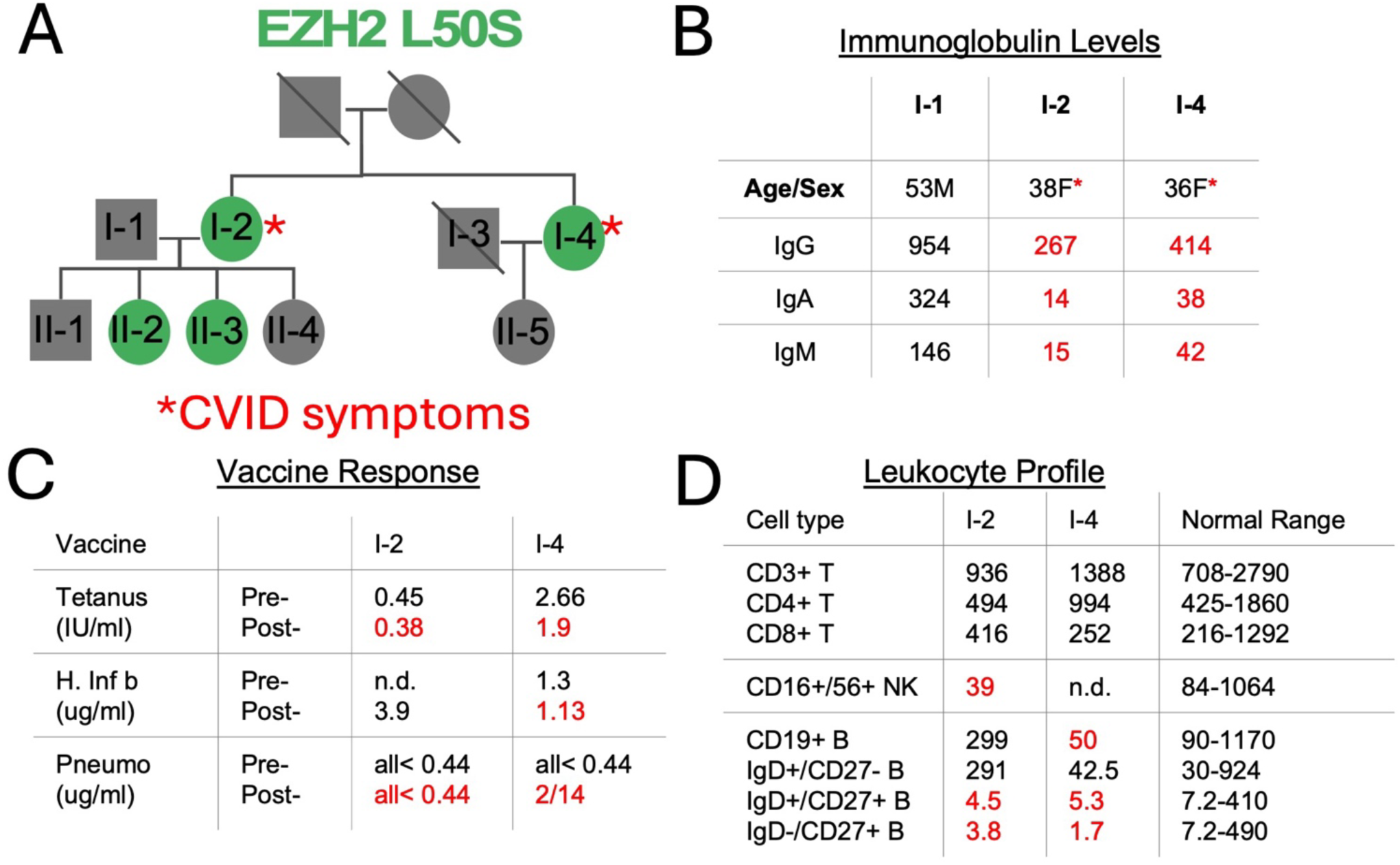
Identification of a variant in EZH2 associated with B cell immunodeficiency. A) Family tree with CVID patients identified, along with EZH2 variant status (heterozygous) indicated by green shading. B) Ig profiling at the time of diagnosis for patients. C) Vaccine challenge antibody titer response, pre- and post-vaccination. Vaccines: *tetanus* toxoid and *H. influenza* b conjugate vaccines and pneumococcal vaccine (Pneumovax-23). D) Flow cytometry profiling of leukocyte type. For B-D, Red indicates below normal levels.

Exome sequencing and subsequent Sanger sequencing was performed within the family cohort, which identified two variants of interest, heterozygous in both sisters, one in the EZH2 gene and one in UNC13D. Certain UNC13D variants are associated with autoimmune lymphoproliferative disease (ALPS)^15^ and hematophagocytic lymphohistiocytosis (HLH). The particular UNC13D variant, Val781Ile, has a high estimated heterozygous frequency in the general population (1 in ∼250 individuals, per gnomAD^16^) and is variably classified in ClinVar as “likely benign” and “uncertain significance”^17^, none of the family members have had clinical disorders that resemble ALPS or HLH, and hypogammaglobulinemia is not generally observed in HLH. The EZH2 variant, NM_004456.5 (EZH2): c.149>C (p. Leu50Ser) has for some time been categorized as ‘likely pathogenic’ based on its association with a patient with Weaver Syndrome^18^, an overgrowth condition frequently caused by mutations in subunits of the Polycomb Repressive Complex 2^19^, of which EZH2 is the catalytic subunit. However, in the midst of our study, this variant has been re-classified as being of ‘conflicting significance’ with the increase in available genome sequences from individuals (see below for more discussion) and lack of any functional studies (prior to the current study). EZH2 is a known driver of B cell development^2,20^ and maturation^4,21^. Based on this, we hypothesized that the L50S variant may have reduced function, resulting in an eventual adult onset of B cell deficiency.

### EZH2 L50S is a rare variant in a conserved protein sequence located in the EED binding domain

Mapping of the L50 position of EZH2 onto the cryo-EM structure of PRC2^22,23^ identified the residue to be in the EED binding domain within a long alpha helix that spans the “back side” of EED, opposite the pocket that binds methylated H3K27 for allosteric stimulation of the complex (Figure 2A,B). Leucine 50 is positioned more on the outward face of the helix, rather than buried in the interface with EED, though this varies subtly in different structures. Sequence alignment of EZH2 protein with vertebrates and *D. melanogaster* revealed that L50 is highly-conserved across nearly all mammals and many vertebrates and in a region with at least three flanking amino acids on either side conserved to zebrafish (Figure 2C, Supplemental Figure 1 for full alignment). However, *D. melanogaster* does not have a similar region. We noted that there are some species where this position differs from leucine, which may help explain the tolerance for L50S.

**Figure 2.**
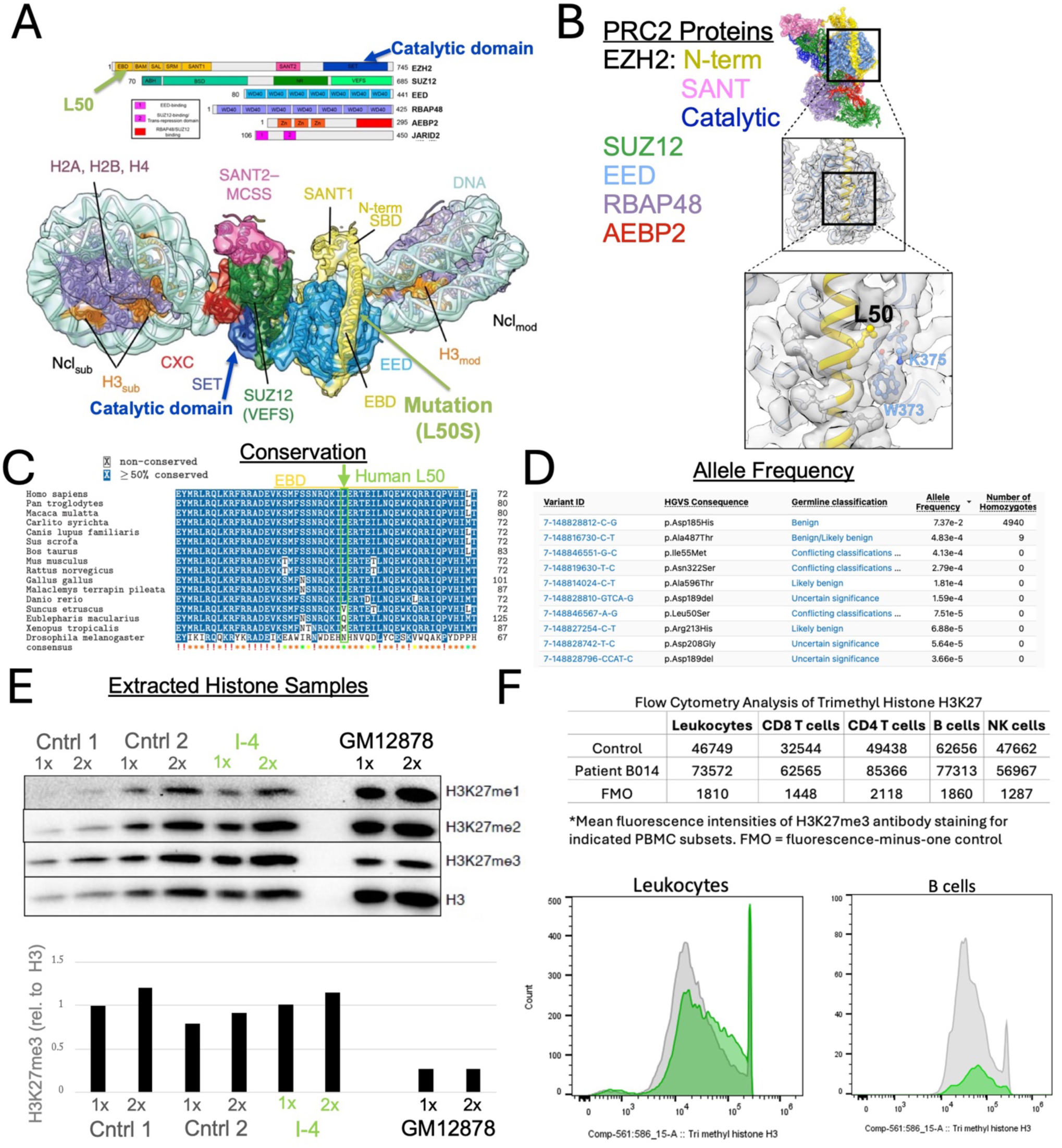
EZH2-L50S structural, genetic, and patient histone sample analysis. A,B) Location of L50 relative to the primary and tertiary structure of PRC2 bound in between two nucleosomes. Adapted from^23,27^. C) EZH2 protein sequence region alignment across vertebrates, highlighting conservation of L50 and divergence in certain species, including non-vertebrate *D. melanogaster*. Consensus symbol key: ! = 16/16; orange * = 15/16; yellow * = 14/16; green * = less than 14/16. D) Data from the gnomAD database, indicating the top 10 most frequent EZH2 alleles, including L50S. E) Extracted histone samples from control or I-4 patient PBMCs or a B cell line(GM12878) analyzed by western for histone H3K27 methylation or total histone H3. F) Leukocyte flowcytometry analysis of Control 2(gray) or I-4 patient(green) to quantify cell types and simultaneously H3K27me3 levels. (*Bottom)* Histogram of all leukocytes (*left*) and B cells (*right*). FMO is a control with all antibodies except for H3K27me3.

With the high conservation among vertebrates, we next asked what the extent of this variation was in the human population. Our initial identification of the L50S association with the patients occurred prior to the large influx of individual exome and genome sequences that was made available through repositories such as gnomAD^16^. With the release of gnomAD v4, greater evidence of L50S in the human population was revealed, suggesting that L50S is a rare, but significant variant, with an allele frequency of ∼7×10^-5^, estimating it in the top 10 most frequent missense alleles of EZH2 present in the human population, with no homozygotes reported. A much lower occurrence of L50V is also reported in the gnomAD database. With an estimated 1 in ∼10,000-20,000 individuals harboring this variant, according to all current UK Biobank^24^ and AllofUs^25^ data, we were motivated to further characterize whether there may be an altered function of this EZH2 L50S variant that has a significant frequency in the human population.

### EZH2 L50S variant blood samples do not show reduced bulk histone methylation

Since EZH2 is a methyltransferase primarily for histone H3 lysine 27, we used peripheral blood mononuclear (PBMC) samples from one of the affected sisters to characterize total H3K27 methylation. Histones were extracted by the high salt method according to standard protocol^26^. Quantification of the three states of histone methylation, normalized to total histone H3, demonstrated no significant loss in histone methylation in the PBMC sample compared to a control hayfever patient (Figure 2E). Additional experiments using flow cytometry with permeabilization to profile H3K27me3 confirmed that no significant change in histone methylation was observed in the patient sample across leukocytes and specifically in the remaining B cells (Figure 2F, Supplementary Figures 2,3). We concluded from this that either the EZH2 variant L50S does not affect histone methylation, or that the context of the patient samples was not optimal to address this. We noted that since B cell maturation requires high EZH2 activity, and the patient samples had abnormally very low B cell count (19 cell/µL at sample collection, years later than in Figure 1, ∼5 times below minimum normal range), that the blood samples may be depleted for the very cell type most affected. Due to these low B cell numbers in the patient blood, it has been challenging to pursue more directed questions of the activity of the variant. In light of this, we wished to establish a system to study the EZH2-L50S variant more directly and with the capacity for a wider series of experimental approaches.

### An inducible system to study EZH2-L50S demonstrates stable variant protein and a shift in histone methylation pattern

To further study the EZH2 L50S variant, we constructed multiple clones of an inducible cell line that expresses wild-type (WT) or L50S EZH2 transgene, introduced at a specific genetic ‘safe harbor’ locus via Cre-mediated recombination^28^. Briefly, HEK 293T cell line was a previously-engineered to include a selectable marker with LoxP sites flanking it^29^. We introduced into this cell line a plasmid carrying a FLAG epitope-tagged EZH2, WT or L50S, and a different selectable marker, flanked by LoxP sites, transfected with a plasmid expressing Cre recombinase and stable cell lines were selected with antibiotic (Figure 3A). We first characterized the expression of EZH2 in these clones and observed reproducible similar expression patterns for WT and L50S EZH2 transgene, which can be distinguished from endogenous EZH2 by size, the transgene is FLAG-tagged, and also with specific western probing for the FLAG epitope (Supplemental Figure 4). Wild-type and L50S EZH2 were both expressed at similar levels, demonstrating that the L50S variant does not destabilize the protein (Figure 3B). Interestingly, we noted that induction of transgene expression led to near complete reduction of the endogenous EZH2, apparently by a post-translational mechanism that we have yet to uncover and do not focus on further in the current study. In effect, due to the transition from endogenous to transgene EZH2 protein, we were able to obtain a ‘pseudo-homozygous’ context to study the EZH2 L50S protein, in the absence of nearly all WT endogenous EZH2.

**Figure 3.**
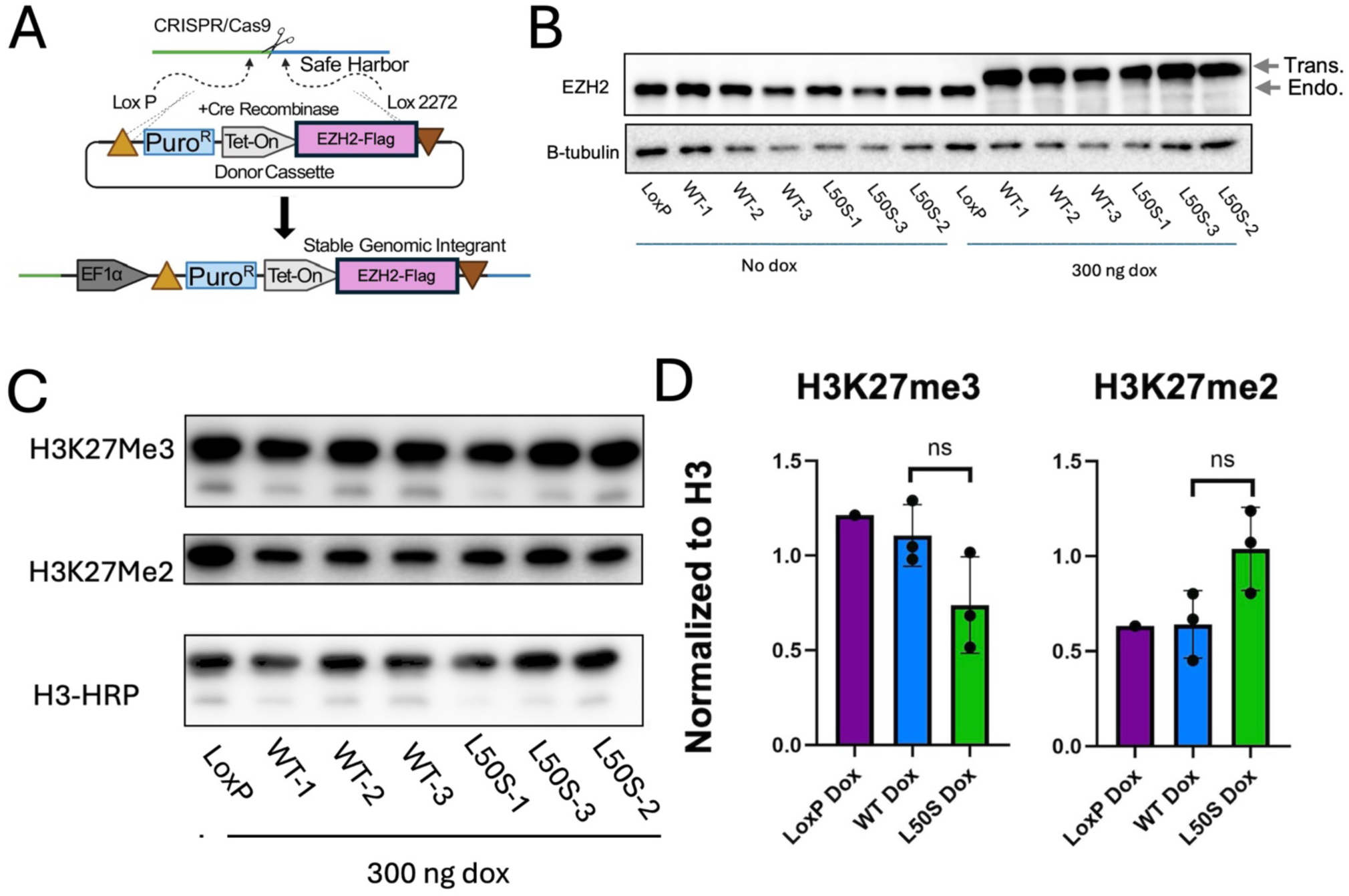
EZH2-L50S is stable and expression leads to only minor H3K27 methylation changes. A) Schematic of the safe harbor locus transgene insertion strategy, adapted from^28^. B) Transgene cell line tests of EZH2 induction via western analysis. Endogenous EZH2 protein (Endo.) and transgene FLAG-EZH2 protein (Trans.) indicated. C,D) Western analysis of histone methylation with EZH2 transgene induction with quantification in D).

We took advantage of the transgene predominance to first determine the levels of H3K27 methylation in WT and L50S contexts. Western analysis of H3K27 di- and tri-methylation demonstrated a shift in signal normalized to the total histone H3, where the L50S transgene generated a lower level of H3K27 trimethylation in most lines, followed by a higher level to H3K27 di-methylation. While there was variation among the clones and the trend was not statistically significant by general p-value standards, this shift in signal suggested that EZH2-L50S may be affecting some small population of sites in the genome, reducing the levels of H3K27 trimethylation.

### CUT&Tag epigenomic profiling reveals regions of H3K27me3 loss in the EZH2 L50S context

To analyze effects of EZH2-L50S compared to wild-type across the genome, we used Cleavage Under Targets and Tagmentation (CUT&Tag)^30^, an antibody-tethered method for probing epigenomic state in situ with isolated whole nuclei. H3K27me3 CUT&Tag was performed with LoxP parental cell line or cell lines with wild-type or L50S EZH2 induced as above (Figure 4A). Libraries from H3K27me3 CUT&Tag samples that were generated and sequenced all demonstrated prominent mononucleosome-sized fragments (Supplemental Figure 5), which were used for further analysis. Combined peaks for WT samples were called and the window around the peak summits analyzed. While all L50S samples demonstrated enrichment of reads within the majority of WT peak regions (Supplemental Figure 6), the overall signal was reduced (Figure 4B). Differential enrichment analysis was performed on the WT versus L50S samples, identifying a small set of peaks that were higher in WT (WT>L50S), while the majority were unchanged (WT=L50S) (Figure 4C, Supplemental Files 1 and 2). We noted that the WT>L50S peaks tended to be those with higher intensity within all peaks, whereas the WT=L50S peaks tended to be average to low intensity (Figure 4D). The fact that no significant WT<L50S peaks were called suggests a high likelihood for L50S to negatively affect H3K27me3 (Figure 4E). We compared the locations of the differential peaks to annotations of gene features including promoter proximity (Figure 4F,G). WT>L50S peaks were highly enriched near promoters, with more than ∼75% overlapping within the window from 7kb up or downstream of a promoter (Figure 4F). Gene Ontology analysis of WT>L50S genes demonstrated enrichment for neuronal biological processes (Supplemental File 3), consistent with neuronal genes normally being repressed in HEK293 cells^31^. Conversely, WT=L50S peaks displayed a distribution pattern similar to shuffling the peaks. WT>L50S peaks were roughly evenly balanced in their propensity to be up or downstream of the TSS with a slight bias towards downstream (Figure 4G). Since the majority of peaks were unchanged, the CUT&Tag analysis helped confirm that the overall H3K27me3 levels are not majorly affected by L50S. We were thus able to highlight specific loci, mainly at highly methylated promoter regions in WT context, that lost significant H3K27me3 in the L50S context.

**Figure 4.**
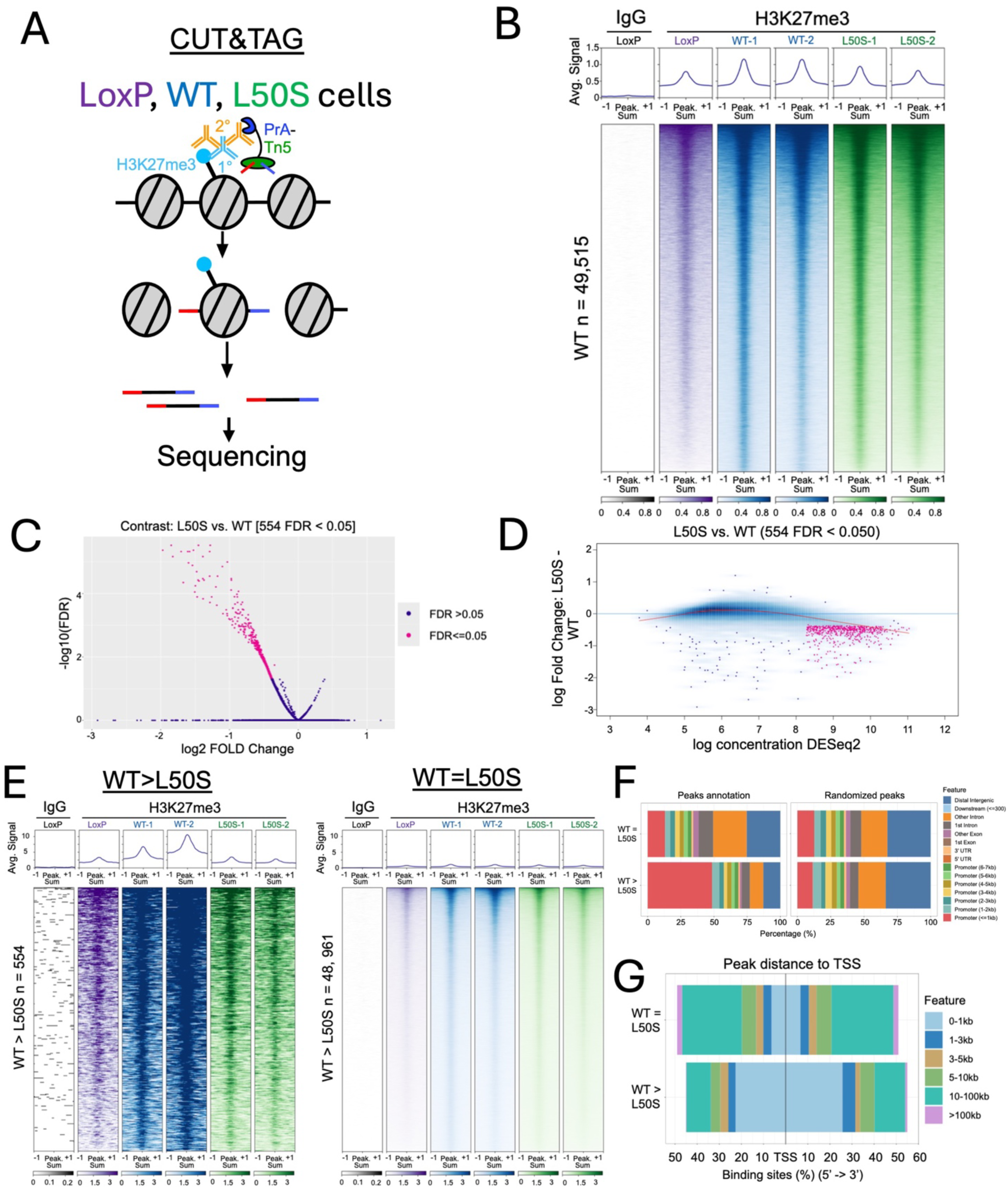
Epigenomic profiling of H3K27me3 in EZH2-L50S expressing cells. A) Schematic for the CUT&Tag experiments performed. B) Heatmaps of CUT&Tag for H3K27me3 (or IgG, far left) highlighting the peak regions identified in the wild-type transgene context, +/- 1 kilobases from peak summit. C) Volcano plot for differential H3K27me3 across the WT and L50S samples and regions in B). Highlighted in magenta are regions that are significantly different between the two types of samples. D) Plot of signal intensity vs. fold-change difference between WT and L50S samples, significant differences in magenta. E) Heatmaps broken out into classes of loci, WT>L50S and WT=L50S. F) Genomic annotation of groups from E) and a set of the same sized windows randomly distributed across the genome as a reference. G) Distance of the peak summits from E,F) to the nearest TSS, indicating relative position, upstream or downstream.

### EZH-L50S is still recruited to promoters yet is deficient at H3K27 methylation there

Since the majority of changes in the WT>L50S H3K27me3 analysis were near promoter regions, we focused in on those for further analysis (Figure 5A). To gain further insight into the mechanism for H3K27me3 loss in those regions, we profiled WT and L50S FLAG-tagged EZH2 by CUT&Tag. We reasoned that either PRC2 with EZH2-L50S was deficient in recruitment to regions of high WT activity or that the L50S complex, once recruited, was intrinsically less active at those regions where WT can operate robustly. Somewhat surprisingly, we found that EZH2-L50S was recruited at a similar, if somewhat higher, level to the promoters with significant WT H3K27me3. Specifically at regions that were significantly lower in H3K27me3 in L50S compared to WT, we observed no decrease, even a slight increase, in recruitment of EZH2-L50S and presumably the full PRC2 complex. To confirm that the FLAG antibody used to identify binding sites of WT or L50S was specific to EZH2, we performed CUT&Tag with an EZH2 antibody in WT cells and observed highly similar enrichment to the FLAG-tagged WT EZH2 that was profiled by FLAG antibody (Figure 5A).

**Figure 5.**
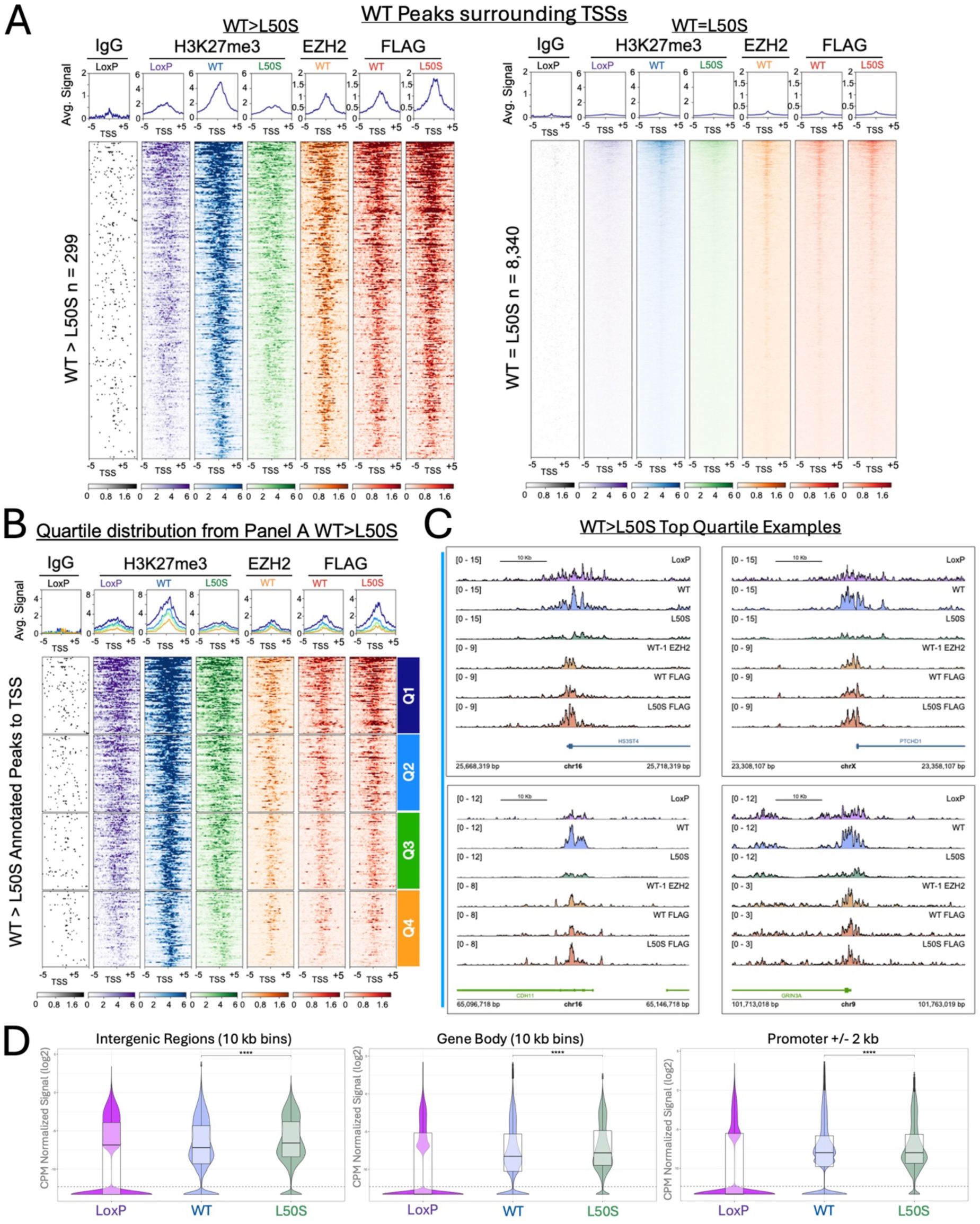
EZH2-L50S is recruited to regions that it cannot properly methylate. A) Heatmaps centered on peaks from Figure 4A that occur within 2 kb of a TSS. Included are combined profiles of H3K27me3 CUT&Tag as well as CUT&Tag for EZH2 and FLAG epitopes, for indicated cell line contexts, +/- 5 kilobases from the TSS. B) Quartile separation of WT>L50S H3K27me3 peaks form A). C) Genome browser tracks from randomly chosen loci from Quartile 1 in B). D) Analysis of H3K27me3 CUT&Tag reads across three classes of genomic region, intergenic, gene body, or promoter proximal. Statistical significance between the indicated samples is shown (Wilcoxon’s test); *\*\*\*\*p*<0.0001.

We separated the WT>L50S promoter windows into quartiles, based on differential H3K27me3 (Figure 5B). In the top quartile, whereas WT H3K27me3 signal is strong directly over the peak of EZH2, L50S H3K27me3 signal is depleted, even as a strong peak of EZH2-L50S itself is observed. Individual examples of this epigenomic profiling, randomly selected from the top quartile of WT>L50S promoter regions are shown (Figure 5C). Profiles of individual loci are representative of the high WT and low L50S H3K27me3 signal and demonstrate EZH2 recruitment in both WT and L50S contexts. Since the overall levels of H3K27me3 in bulk are not statistically different in L50S, we asked whether there was a compensation of methylation outside of the differential regions. To do this we analyzed overall read counts in three categories that cover all genomic coordinates: intergenic, gene body, and promoter-proximal regions (Figure 5D). We observed higher signal for L50S in intergenic and gene body regions than WT, which may explain overall levels of H3K27me3 being only mildly affected by the variant. This result is consistent with a higher proportion of unique L50S H3K27me3 peaks than WT peaks and more EZH2-L50S peaks outside of the shared WT/L50S peaks (Supplemental Figure 6).

### EZH2-L50S assembles into PRC2 but is deficient in histone methylation activity toward nucleosomes

After identifying a deficiency in the ability of EZH2-L50S to perform histone H3 lysine 27 tri-methylation at regions of high wild-type activity, we next wished to further characterize why this EZH2 variant has lower activity in cells. We first tested whether the L50S mutation in EZH2 disrupted a key subunit interaction within the core of PRC2. While we had already determined that EZH2-L50S could be recruited to PRC2 targets, equally to the WT subunit (Figure 5A), it was still possible that the reduced activity after recruitment was due to a key core subunit that was missing from the L50S complex. We performed co-immunoprecipitation with the FLAG antibody for either WT or L50S EZH2 and probed by western blotting for the core PRC2 subunits SUZ12 and EED. In both cases, both WT and L50S associated with an equal amount of each subunit (Figure 6A), suggesting that the EZH2-L50S variant does not disrupt core PRC2 interactions. This is consistent with the normal recruitment profile for EZH2-L50S (Figure 5A), since SUZ12 is crucial for EZH2 recruitment^32^. The fact that EED can still associate with EZH2-L50S is important, since the L50 position lies in the EED-binding domain helix of EZH2 (Figure 2A,B). Our results demonstrate that the L50S variant does not lead to a major loss of interaction with the core PRC2 subunits.

**Figure 6.**
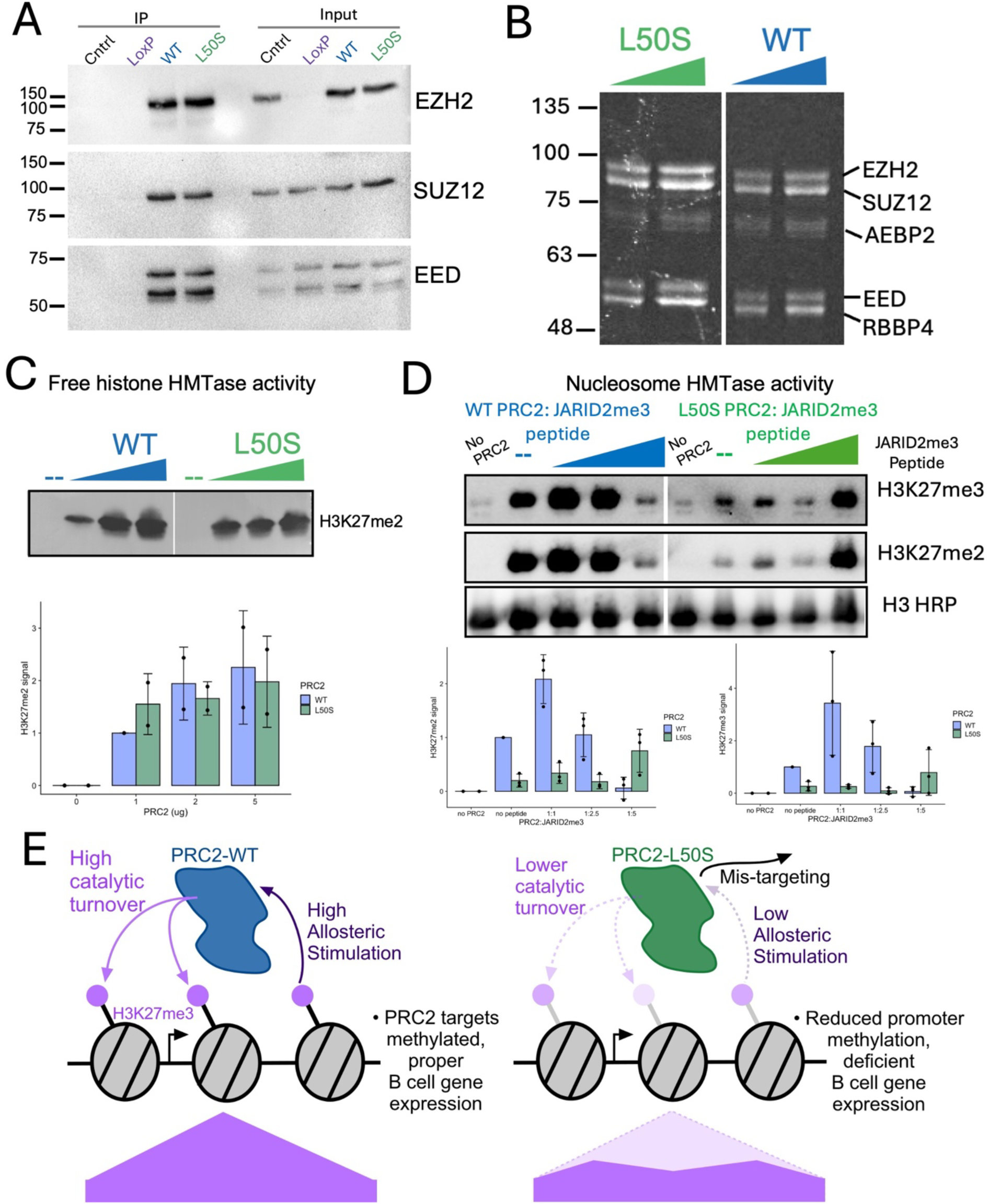
EZH2-L50S leads to lower PRC2 activity and allosteric stimulation on nucleosomes. A) Co-immunoprecipitation experiments using anti-FLAG antibody, with indicated cell lines. Western analysis performed for EZH2, SUZ12, and EED. Control indicates identical IP conditions with no FLAG antibody using the WT cell line. B) SDS-PAGE and Coomassie-stained, recombinant, baculovirus-expressed and purified five-subunit PRC2 complex with EZH2-WT or -L50S. C) Histone methyltransferase (HMTase) assays with recombinant PRC2 using free histone samples. H3K27me2 western and quantification shown. D) Histone methyltransferase (HMTase) assays with recombinant PRC2 using a nucleosome substrate with increasing titration of a stimulatory methylated peptide corresponding to a region of JARID2. E) Model of EZH2-L50S effect on promoter-proximal H3K27 methylation. The variant lowers PRC2 catalytic turnover and dynamics of the complex at promoter chromatin, in part by lowering the ability of allosteric stimulation by H3K27 methylation itself. Flanking chromatin, methylated by PRC2.2, is less affected. PRC2-L50S also exhibits some mis-targeting away from promoter regions, where it displays a diffuse methylation pattern.

To continue to directly characterize the EZH2-L50S variant in PRC2, we used an established baculovirus expression and purification system^33^ to isolate purified human PRC2. Subunit stoichiometry of the L50S version was identical to WT (Figure 6B). We first tested the general histone methylation ability of the purified complex with pure recombinant histones, lacking any modifications, a sample of human histone octamer that exists as H3/H4 tetramers and H2A/H2B dimers under the PRC2 reaction conditions. Under these conditions, we observed no significant difference in activity between WT and L50S complexes (Figure 6C). Next, we tested the methylation activity on nucleosomes, a more complex substrate that, through its assembly, requires a specific context to promote histone methylation, especially di- and tri-methylation of H3K27^34^. Incubation of L50S with a nucleosome containing a linker DNA sequence produced a lower amount of H3K27 methylation (both di- and tri-methylation), roughly 25% compared to WT PRC2 (Figure 6D, “No peptide”). We next asked whether the PRC2 activity that is stimulated allosterically by binding of the EED subunit to a methylated peptide may be affected in the L50S context, since leucine 50 is in the EED-binding domain of EZH2. We titrated a synthetic trimethylated peptide into the nucleosome histone methyltransferase assay and observed a deficiency in the ability of PRC2-L50S to allosterically respond (Figure 6D).

Our results identify a hypomorphic EZH2 variant, L50S, in a family with B cell immunodeficiency. PRC2-L50S can assemble and bind promoters, but its activity is weak at promoter regions where wild-type PRC2 is most active, whereas L50S has higher propensity to methylate in a distributive manner in non-promoter regions (Figure 6E). PRC2-L50S is less active on nucleosomes and does not respond normally to allosteric stimulation that is mediated through EED.

## Discussion

While EZH2 expression has been known for some time to be induced during B cell activation and EZH2 knockout disrupts B cell function during an immune response^4,21^, our work connects a human genetic context in a B cell immunodeficiency with a hypomorphic variant in EZH2. The adult-onset loss of B cells is likely buffered earlier in life by the wild-type EZH2 allele and the fact that L50S is not a complete loss of function or a dominant negative, nor does it directly affect the PRC2 active site. In fact, our examination of overall histone methylation levels detected only minor differences and only in the transgene context (Figures 2 and 3). This same observation is seen in PRC2 accessory protein knockouts, where even the triple knockout of all PCLs, JARID2 and AEBP2, which drastically affects H3K27me3 at PRC2 targets, has minimal effect on bulk H3K27 methylation^35^. Despite a general maintenance of overall histone methylation, we see explicit deficiency of L50S to support wild-type level H3K27me3 at primarily promoter regions where wild-type PRC2 activity is very high (Figure 5A). Surprisingly, EZH2-L50S is recruited to a higher level than wild-type to these affected regions, suggesting that L50S leads to PRC2 getting “stuck”, but low in activity. Our biochemical analysis demonstrates a loss of nucleosome methylation and EED allosteric stimulation, suggesting that this could lead to improper B cell gene expression (Figure 6E). Overall, this work sheds new light on the regulation or PRC2 and highlights a new player in the population genetics of immunity.

The pattern of H3K27me3 at promoters that are deficient for methylation in the L50S context is reminiscent of H3K27me3 in a knockout for all PRC2.1 activity (PCL 1-3, triple KO), observed in previous work^35,36^, with a trough observed where the peak intensity is seen in WT. In this background, PRC2.2 remains to maintain the “arms” of the methylation peak. PRC2.2 is distinguished by incorporation of AEBP2 and JARID2 accessory factors. The methylated peptide used in our characterization of purified EZH2-L50S (Figure 6D) is derived from the region of JARID2 that is normally methylated by EZH2 and which can promote allosteric stimulation of EZH2 when JARID2 associates in PRC2.2^37^. This mechanism operates in a similar manner to allosteric stimulation when the histone H3K27me3 tail binds to EED^8^. While the allosteric stimulation of the peptide, provided in trans, is affected in the L50S context, at high concentrations, the peptide begins to stimulate PRC2-L50S (Figure 6D). When full-length JARID2 forms a stable complex with PRC2, this peptide is likely present at high local concentration to provide some level of allosteric stimulation, which may help explain why the H3K27me3 pattern in the L50S context appears similar to the pattern when PRC2.2 is active, but PRC2.1 is not. The accessory factors for PRC2.1, the PCLs, are DNA binding proteins which are thought to help recruit/stabilize PRC2.1 to the regions that gain highest H3K27me3^35^. Despite the histone methylation pattern similarity to PCL KO, we do not observe a loss of EZH2 recruitment, and presumably the full PRC2 is bound even at the regions that lose the most H3K27me3. This behavior of normal/higher PRC2 recruitment, yet lower H3K27 methylation, more closely resembles mutants that disrupt the protein interaction network that promotes allosteric stimulation from EED methylated peptide binding to higher EZH2 activity^10^. Currently, we interpret the L50S context as potentially a combination effect on PRC2.1 mechanism and disruption of allosteric activation, where both could contribute to the molecular outcome. This suggests a de-coupling of the PCL-dependent mechanism, favoring JARID2-dependent activity. Interestingly, JARID2 was found to be down-regulated by a microRNA in B cells during germinal center formation^38^, which prevents JARID2-mediated apoptosis. We predict that EZH2-L50S may tip the balance toward JARID2, which may then lead to B cell apoptosis instead of expansion in germinal centers. This would help explain how EZH2 is required for B cell expansion in GCs^4^, perhaps through a PRC2.1-dominant mechanism. This finding is consistent with a role for PCL3 (PHF19) from PRC2.1 in GC B cell expansion and production of antibody-secreting cells^39^. This model is also supported by recent work demonstrating that loss of JARID2 promotes B cell output while loss of EZH2 reduces B cell differentiation and exit from the bone marrow^40^.

In conclusion, our work highlights a human genetic context associated with B cell dysfunction and loss, resulting in CVID, based on a hypomorphic variant that affects the Polycomb system. The CVID patients developed CVID in adulthood, suggesting that EZH2-L50S, as a heterozygous genetic context, does not affect human development as Weaver Syndrome does. This is likely due to the compensation of the wild-type allele and the fact that EZH2-L50S is not as severe a loss of function as Weaver Syndrome mutations, nor does it act as a dominant negative as WS mutations can^41^. With the significant prevalence of this rare variant in the human population, inclusion of EZH2 testing in immunodeficiencies should be considered, and future work is needed to determine if other factors may interact to determine the clinical outcome of those that bear EZH2-L50S.

## Materials and Methods

### Patient testing in the clinic

The clinical samples were collected according to institutional guidelines and with the approval of the Colorado Multiple Institutional Review Board. Profiles for IgG, IgM and IgA and were measured by standard clinical laboratory methods. Immunization tests were done with commercially available, FDA-approved vaccines. Pneumovax-23 (Merck), Hemophilus b conjugate (GSK) and tetanus toxoid (& Diptheria) (Sanofi Pasteur) were given in the standard doses. Serum for antibody titers and measurements of antibody avidity were collected prior to immunization and approximately 4 weeks after immunizations. These vaccines were selected because the antibody response to 1) (tetanus) is T-cell dependent; 2) (pnuemovax-23) is T-cell independent; and 3) (hemophilus b conjugate) depends on T-cell help. After the blood was clotted, serum was collected from centrifuged samples and stored at -80^°^C. Titers of antibodies in serum pairs were assayed together using commercial multiplex bead assays as specified by the manufacturers. Expected “normal” responses for the purposes of this study, as suggested in the Practice Parameter for the Diagnosis and Management of Primary Immunodeficiency^42^, are ≥ 4-fold increases in antibody titers after immunizations with tetanus toxoid and Hemophilus influenza b conjugate. Expected results after immunization with 23-valent pneumococcal vaccine are either a ≥4-fold increase in titer or attainment of a titer of 1.3 µg/ml or greater for at least 70% of the serotypes that are assayed^42^. Lymphocyte populations and sub-populations were quantitated from heparinized venous blood that had been centrifuged on Ficoll-Paque gradients. The mononuclear cells were tagged with selected monoclonal antibodies and quantitated by flow cytometry.

### Patient genetic sequencing

The clinical samples were collected according to institutional guidelines and with the approval of the Colorado Multiple Institutional Review Board. Patient PBMC samples from Ficoll-Paque gradient were subjected to standard DNA extraction protocols and prepared for Illumina sequencing library. Whole genome sequencing at 125X coverage was performed by the University of Colorado Comprehensive Cancer Center Genomics Core Facility.

The freeze 9 genotype call set is produced by the Freeze 8 variant calling pipeline performed by the TOPMed Informatics Research Center (Center for Statistical Genetics, University of Michigan, Hyun Min Kang, Jonathon LeFaive and Gonçalo Abecasis). The software tools in this version of the pipeline are available on github at https://github.com/statgen/topmed_variant_calling. The following description refers to specific components of the pipeline. These variant calling software tools are under continuous development; updated versions can be accessed at http://github.com/atks/vt or http://github.com/hyunminkang/apigenome.

The GotCloud pipeline detects variant sites and calls genotypes for a list of samples with aligned sequence reads. Specifically, the pipeline for freeze 8 consists of the following six key steps: Sample quality control, Variant detection, Variant consolidation, Genotype and feature collection, Inference of nuclear pedigree, Variant filtering. These procedures have been integrated into the current release of the GotCloud software package at https://github.com/statgen/topmed_variant_calling.

### Flow Cytometry

Cryopreserved PBMC were thawed in 1 x Phosphate Buffered Saline (PBS) containing 1% Fetal Bovine Serum (FBS) and 0.1% Sodium Azide and blocked for 10 min at room temperature in 1 x PBS containing 10% human serum and 10% mouse serum. After washing, cells were incubated for 30 min at 4°C with antibodies specific for CD3, CD4, CD8, CD11b, CD11c, CD14, CD15, CD16, CD19, CD45, CD56, CD66, and HLADR. Cells were washed and permeabilized with Transcription Factor Buffer Set (BD Pharmingen) and incubated for 1 h at 4°C with antibodies specific for tri methyl histone H3. Samples were collected on an LSR Fortessa X-20 Cell Analyzer (BD Biosciences) and analyzed with FlowJo v10 software.

### Protein alignment

EZH2 orthologs protein sequences from the indicated species were obtained from the NCBI database. Sequence alignment was performed using msa package in R^43^ that provides an interface for Clustal W^44^.

### Plasmids

Wild-type and L50S EZH2 coding sequences were cloned into pRD-RIPE ^29^ to be used in Cre/Lox generation of cell lines (see below).

### Cell culture and cell lines

Parental LoxP HEK293T^29^ and derived cell lines were routinely grown in Dulbecco’s Modification of Eagle’s Medium 1x with 4.5 g/L glucose, L-glutamine and sodium pyruvate (Corning, DMEM1x). DMEM 1x was supplemented with 10% (v/v) Fetal Bovine Serum (FBS, Avantor Seradigm), 100 IU/mL Penicillin/ and 100 µg/mL Streptomycin (Gibco). Blasticidin (Gibco) 300 ng/mL and Puromycin 3 µg/mL (Sigma-Aldrich) were added to the growth media when needed.

All LoxP-containing cell lines followed the same protocol for cassette switching, as described^28^. Cells were co-transfected with cassette plasmid with 1% (wt/wt) of a Cre encoding plasmid (pBT140, addgene #27493). Donor plasmids with transgenes have a tetracycline transactivator construct, as well. Plasmids were transfected with Lipofectamine LTX reagent (Thermo Fisher Scientific 5338030) and Opti-MEM (Thermo Fisher Scientific 31985070) following the manufacturer’s protocol for 24 hr. After 24 hr in non-selective medium, puromycin-resistant colonies were expanded in growth medium supplemented with 5 μg/mL puromycin. Three independent WT and L50S cell lines were produced through this method (WT-1, -2, 3; L50S-1, -2, -3).

### Immunoblotting

Cells were grown as described above in 10-cm dishes, washed once with 1X PBS, treated with 1 ml of 0.25% trypsin (Corning), pelleted, and supernatant was discarded. Approximately 1 x 10^6^ cells were then homogenized as described^45^ with 200 µL of TOPEX+ buffer (50mM Tris-HCl pH7.5,300 mM NaCl, 0.5% Triton X-100, 1% SDS, 1mM DTT, 1X Complete EDTA-free Protease Inhibitor (Roche Applied Science), 33.33 U/mL Benzonase Nuclease HC (EMD-Millipore). Samples were then incubated for 20 min on ice, and then centrifuged for 20 min, 4°C at 13,000 rpm. Supernatant was transferred to a fresh tube, and protein concentration was determined using the Pierce BCA Protein Assay Kit (Thermo-Scientific). Samples were run on SDS-PAGE gels, transferred to nitrocellulose or PVDF membranes, and blocked with TBST (50 mM Tris pH 7.5, 150 mM NaCl, 0.1% Tween-20) containing 5% milk for 30 min at room temperature. Membranes were then incubated in TBST with 3% BSA containing one of the following primary antibodies: H3K27me3 (9733, Cell Signaling Technology) with a dilution of 1:5,000 during 1 hour at room temperature; H3K27me2 (9728, Cell Signaling Technology) with a dilution of 1:5,000 during overnight at 4°C; H3K27me1 (61016, Active Motif) with a dilution of 1:1,000 during overnight at 4°C; H3-HRP (21054, Abcam) with a dilution of 1:100,000 during 1 hour at room temperature; EZH2 (5246, Cell Signaling Technology) with a dilution of 1:2,000 during overnight at 4°C; FLAG (F1804, Sigma-Aldrich) with a dilution of 1:400 during overnight at 4°C; SUZ12 (D39F6, Cell Signaling Technology) with a dilution of 1:500 during overnight at 4°C; EED (85322, Cell Signaling Technology) with a dilution of 1:1000 during overnight at 4°C; B-tubulin (Y1061, UBPBio) with a dilution of 1:1,000 during overnight at 4°C. Membranes were then washed 3 times for 5 min and incubated in TBST with 3% BSA containing the corresponding anti-rabbit or anti-mouse secondary HRP-conjugated antibody (1706516 or 1705046, BIORAD) with a dilution of 1:20,000 during 1 hour at room temperature. Membranes were then washed 3 times for 5 min with TBST and visualized using chemiluminescence Western Chemiluminescent HRP Substrate (WBKLS0500, Immobilon).

### CUT&Tag

H3K27me3, EZH2, and FLAG signals were profiled using the CUTANA Direct-to-PCR CUT&Tag Protocol (V2) from EpiCypher. Briefly, cells were grown under doxycycline inducible conditions (300 ng/mL) for 8 days, harvested with 0.25% Trypsin-EDTA solution (Corning, 25-053-cl) and counted in Countess Cell Counting Chamber Slides (Invitrogen, C10228) using 0.4% Trypan Blue Solution (Gibco, 15252061).

To extract nuclei, a total of 1 x 10^6^ cells were washed in 1 mL 1x PBS, then 1.1 x 10^5^ cells were centrifuged and resuspended in 100 µL cold Nuclei Extraction Buffer (20 mM HEPES-KOH pH 7.9, 10 mM KCl, 0.1% Triton x-100, 20% glycerol, 0.5 mM spermidine and 1x Complete protease inhibitor(Millipore-Sigma). To bind nuclei to CUTANA Concanavalin A Beads (EpiCypher, 21-1401), nuclei samples were centrifuged, resuspended in 105 µL of cold Nuclei Extraction Buffer, and 100 µL of sample were transferred into a 200 µL tube containing 10 µL of activated Concanavalin A Beads. To bind primary antibody to target regions, bound nuclei on Concanavalin A Bead samples were centrifuged, resuspended in 50 µL of cold Antibody 150 Buffer (20 mM HEPES-KOH pH 7.5, 150 mM NaCl, 0.5 mM spermidine, 1x protease inhibitor, 0.01% digitonin, 2 mM EDTA), then specific antibodies were added to the samples and incubated overnight at 4 °C. Antibodies used: 1 µL of H3K27me3 (9733, Cell Signaling Technology), 1 µL of EZH2 (5246, Cell Signaling), 1 µL of anti-FLAG (F1804, Sigma-Aldrich) or 0.5 µL IgG (13-0042, Epicypher).

To bind secondary antibody to primary antibody, 0.5 µL of matching secondary antibody, anti-rabbit (13-0047, Epicypher) or anti-mouse (13-0048, Epicypher), were diluted in 50 µL of Digitonin 150 Buffer (20 mM HEPES-KOH pH 7.5, 150 mM NaCl, 0.5 mM spermidine, 1x protease inhibitor, 0.01% digitonin), added to the corresponding primary antibody, and samples were then incubated at room temperature for 30 min. To bind CUTANA pAG-Tn5 (EpiCypher, 15-1017) to secondary antibody, samples were washed two times with 200 µL of cold Digitonin 150 Buffer, resuspended in 50 µL of cold Digitonin 300 Buffer (20 mM HEPES-KOH pH 7.5, 300 mM NaCl, 0.5 mM spermidine, 1x protease inhibitor and 0.01% digitonin), 2.5 µL of pAG-Tn5 was added to each sample, samples were then incubated for 1 hr at room temperature.

To perform Tn5 tagmentation over chromatin, samples were washed two times with 200 uL of cold Digitonin 300 Buffer, resuspended in 50 µL of Tagmentation Reaction Buffer (20 mM HEPES pH 7.5, 300 mM NaCl, 0.5 mM spermidine, 1x protease inhibitor, 10 mM MgCl2, and 0.01% digitonin), and incubated for 1 hr at 37 °C. To stop the tagmentation reaction and perform release of tagmented chromatin into solution, samples were placed on a magnet and supernatant was discarded. Tagmented chromatin bound to magnetic beads was then resuspended in 50 µL of equilibrated to room temperature Pre-Wash 150 Buffer (20 mM HEPES-KOH pH 7.5, 150 mM NaCl). Magnetic beads were placed back on a magnet, supernatant was discarded, and resuspended in 5 µL SDS Release Buffer (10 mM TAPS pH 8.5 and 0.1% SDS). Tagmented chromatin was released by incubation for 1 hr at 58°C and quenched using 15 uL of SDS Quench Buffer (0.67% Triton x-100).

To perform library amplification, 2 µL of i5 universal primer (EpiCypher), 2 µL i7 barcoded primer (EpiCypher) and 25 µL NEBNext High Fidelity 2x PCR Master Mix (NEB, M0541S) were added to tagmented chromatin samples, the following PCR conditions were used: 5 min at 58°C, 5 min at 72°C, 45 sec at 98°C, 16 – 19 cycles of 15 sec at 98°C, 10 sec at 60°C and final extension of 1 min at 72°C. Remaining oligos and undesired PCR products were cleaned up using 1.3X High Prep (MAGBIO, AC60005) magnetic beads following manufacturer recommendations. Amplified libraries were washed away from magnetic beads in nuclease free water. To check for library concentration and fragment size distribution, Qubit 1X dsDNA High Sensitivity Assay Kit (Invitrogen, Q33231) and High Sensitivity D1000 ScreenTape Assay for TapeStation (Agilent, 5067-5584) were used. Libraries were paired-end sequenced by Novogene or University of Colorado Cancer Center Genomics Core services.

### CUT&Tag data analysis

Data analysis was performed using certain approaches described previously^46^. Initial data pre-processing was done using Cutadapt (Martin, 2011): Illumina adaptors were removed from sequencing reads, the reads were then trimmed to a maximum of 140 bp, and shorter reads than 35 bp were discarded.

Trimmed Read 1 and Read 2 alignment to human genome version hg38 was done using Bowtie2^47^ with the following parameters, *--local --very-sensitive-local --no-unal –no-mixed --no-discordant --I 10 --X 2000*. Duplicated reads were removed using the following Samtools^48^ commands, *samtools sort -n <FILE.sam>* | *samtools fixmate -m - -* | *samtools sort -* | *samtools markdup -s -r - <FILE.out.sam>*. Aligned reads in sam format were processed to the bam format using the following commands, *samtools view -b -F 0×04 <FILE.out.sam> > <FILE.out.bam>*. Read length size distribution was analyzed using the command *bamPEFragmentSize* from deepTools2^49^, then fragments within a size from 100 bp to 300 bp were sieved using *alignmentSieve*, and used in the following analysis.

Biological and/or technical sieved replicates were merged using the following command *samtools merge -n <FILE.merged.out.sieved.bam> <FILE1.out.sieved.bam> <FILE2.out.sieved.bam>*. CUT&Tag peaks were called using SEACR^50^ with the following arguments *-c 0.05 -n norm -m stringent*. Bigwig files were generated using *bamCoverage*^49^ with the following arguments *--normalizeUsing CPM --effectiveGenomeSize 2913022398 --extendReads –binSize 10 --blackListFilename <BLACKLIST.file>*. Differential peak analysis was done using DiffBind (http://bioconductor.org)^51^. Peak summits were re-centered and re-sized to +/- 1 Kb using *dba.counts(summits = 2000),* blacklisted regions were excluded from the analysis using *dba.blacklist()*, data was normalized to library size using the argument *DBA_NORM_LIB*, and only reads in peaks were used for the differential binding calculation using the argument *DBA_LIBSIZE_PEAKREADS*. To generate a differential analysis report of all re-centered binding sites, the command *dba.report()* was used. MA plots and volcano plots were generated using the commands *dba.plotMA()* and *dba.plotVolcano()*, respectively. “WT>L50S” peaks with a FDR < 0.05, as well as WT=L50S” peaks were filtered out from the differential analysis report, then annotated to Gencode annotation v47^52^ using ChIPseeker^53^. No “WT<L50S” peaks with a FDR < 0.05 were identified. Annotation bar plot and distance to TSS plot were done using *plotAnnoBar()* and *plotDistToTSS()* functions, respectively. Annotated peaks within +/- 2 Kb to TSSs were filtered, gene coordinates were obtained and the corresponding bed files were generated using homemade R scripts. Heatmaps for scored regions under re-centered peak summits +/- 5 Kb as well as TSS associated peaks were generated using *plotHeatmap* function from deeptools^49^. Genome browser visualization plots were done using *Plotgardener*^54^.

### Code availability

Code available upon request.

### Data availability

GEO submission availability forthcoming.

### Immunoprecipitation

10 μg of FLAG antibody (F1804, Sigma-Aldrich) was pre-bound to 200 μL Dynabeads Protein G (Thermo Fisher Scientific, 10004D) and resuspended in Lysis Buffer (50 mM Tris pH 7.4, 150 mM NaCl, 1% NP-40, 2 mM EDTA pH 8.0 and 1X Complete Protease Inhibitor from Millipore-Sigma). To extract protein complexes, trypsinized cell samples were washed twice with 1X PBS, centrifuged, and proteins were extracted with 1.5 volume more than cell pellet of Lysis Buffer. Protein extracts were then incubated at 4°C for 60 min with rotation and centrifuged at 13,000 rpms in a microfuge for 15 min at 4°C. Immunoprecipitation was performed by mixing 1 mg/ml of protein extract diluted in Lysis Buffer with 200 μL of bead/FLAG antibody suspension and incubated overnight at 4°C with rotation. Immunoprecipitated proteins were then washed three times with Lysis Buffer on a magnet and resuspended in 50 μl 1X SDS buffer. SDS-PAGE and immunoblotting was performed as described above.

### Histone methyltransferase assays

HMTase assays were performed in a total volume of 20 μL containing methyltransferase buffer (50 mM Tris (pH 8.8), 2 mM MgCl_2_, 20% Triton-X, and 1 mM TCEP) with 100 μM *S*-adenosylmethionine (New England Biolabs), 914 nM histone octamer, and recombinant human WT and L50S PRC2 complexes titrated at points of 0, 150, 310, and 770 nM. The reaction mixture was incubated for 30 min at room temperature and stopped by adding 5 μL of 5X SDS loading dye containing BME. After HMT reactions, samples were incubated for 5 min at 95°C and separated on SDS-PAGE gels. Gels were then subjected to wet transfer (30% MeOH transfer buffer) of histones to 0.45-μm polyvinylidene difluoride membranes (Millipore), and protein was detected by Western blot analysis using primary αRb H3K27me2 antibody (Cell Signaling Technology, 9728S, D18C8), secondary antibody (Bio-Rad, #1705046), and H3-HRP (horseradish peroxidase) (Abcam, #ab21054).

HMTase assays were performed in a reaction buffer containing methyltransferase buffer (10mM HEPES NaOH (pH 7.5), 2.5 mM MgCl_2_, 4% glycerol, 0.25 mM EDTA and 0.1 mM DTT). This was used to bring the final reaction volume up to 15 μL, with 99.3 μM *S*-adenosylmethionine (New England Biolabs), 601 nM JARID2, a titration of JARID2me3 peptide from 648nM to 64.9uM, 160 nM nucleosome, and 570 nM recombinant human PRC2 complexes under the following conditions. Nucleosomes were reconstituted according to^55^. The DNA template is a version of the 601 DNA sequence with 38 bp upstream linker and a small downstream 11 bp linker with a sticky end, 196 bp total. The reaction mixture was incubated for 30 min at 30°C and stopped by adding 3.75 μl of 5X SDS loading dye. After HMTase reactions, samples were incubated for 5 min at 95°C and separated by SDS-PAGE. Gels were then subjected to wet transfer (30% MeOH transfer buffer) of histones to 0.45-μm polyvinylidene difluoride membranes (Millipore), and protein was detected by western blot analysis using primary αRb H3K27me3 antibody (Cell Signaling Technology, 9733S, C36B11), H3K27me2 antibody (Cell Signaling Technology, 9728S, D18C8), secondary antibody (Bio-Rad, #1705046), and H3-HRP (horseradish peroxidase) (Abcam, #ab21054).

## Acknowledgements

This work was supported by NIH grant R35GM144358 (A.M.J). The Human Immune Monitoring Shared Resource (RRID:SCR_021985) and the University of Colorado Cancer Center Genomics and Cell Technologies Cores are supported by the NCI University of Colorado Cancer Center (P30CA046934). We thank Srinivas Ramachandran, Giovana Breda Veronezi, and Abby Trouth for helpful guidance and discussions for performing CUT&Tag. We thank Liqun Luo lab for Cre plasmid, Tom Cech lab for the wild-type PRC2 baculovirus constructs, Eugene Makeyev lab for pRD-RIPE and LoxP HEK293T cells. We thank Vignesh Kasinath for help with figure making. We thank Srinivas Ramachandran and Samuel Gonzalez for feedback on the manuscript. This work is dedicated to the memory of Dr. Charles Kirkpatrick, whose dedication to his patients and love of science drove this study forward.

## References

1. Blackledge, N.P. & Klose, R.J., The molecular principles of gene regulation by Polycomb repressive complexes. Nat Rev Mol Cell Biol 22 (12), 815–833 (2021).

2. Su, I.H. et al., Ezh2 controls B cell development through histone H3 methylation and Igh rearrangement. Nat Immunol 4 (2), 124–131 (2003).

3. Beguelin, W. et al., EZH2 enables germinal centre formation through epigenetic silencing of CDKN1A and an Rb-E2F1 feedback loop. Nat Commun 8 (1), 877 (2017).

4. Beguelin, W. et al., EZH2 is required for germinal center formation and somatic EZH2 mutations promote lymphoid transformation. Cancer Cell 23 (5), 677–692 (2013).

5. Guo, M. et al., EZH2 Represses the B Cell Transcriptional Program and Regulates Antibody-Secreting Cell Metabolism and Antibody Production. J Immunol 200 (3), 1039–1052 (2018).

6. Wiggins, K.J. et al., EZH2 coordinates memory B-cell programming and recall responses. J Immunol 214 (5), 947–957 (2025).

7. Yap, D.B. et al., Somatic mutations at EZH2 Y641 act dominantly through a mechanism of selectively altered PRC2 catalytic activity, to increase H3K27 trimethylation. Blood 117 (8), 2451–2459 (2011).

8. Margueron, R. et al., Role of the polycomb protein EED in the propagation of repressive histone marks. Nature 461 (7265), 762–767 (2009).

9. Glancy, E., Ciferri, C., & Bracken, A.P., Structural basis for PRC2 engagement with chromatin. Curr Opin Struct Biol 67, 135–144 (2021).

10. Lee, C.H. et al., Allosteric Activation Dictates PRC2 Activity Independent of Its Recruitment to Chromatin. Mol Cell 70 (3), 422–434 e426 (2018).

11. Oksuz, O. et al., Capturing the Onset of PRC2-Mediated Repressive Domain Formation. Mol Cell 70 (6), 1149–1162 e1145 (2018).

12. Cohen, A.S. et al., Weaver Syndrome-Associated EZH2 Protein Variants Show Impaired Histone Methyltransferase Function In Vitro. Hum Mutat 37 (3), 301–307 (2016).

13. Cohen, A.S. et al., A novel mutation in EED associated with overgrowth. J Hum Genet 60 (6), 339–342 (2015).

14. Deevy, O. & Bracken, A.P., PRC2 functions in development and congenital disorders. Development 146 (19) (2019).

15. Arico, M. et al., Variations of the UNC13D gene in patients with autoimmune lymphoproliferative syndrome. PLoS One 8 (7), e68045 (2013).

16. Chen, S. et al., A genomic mutational constraint map using variation in 76,156 human genomes. Nature 625 (7993), 92–100 (2024).

17. Landrum, M.J. et al., ClinVar: improving access to variant interpretations and supporting evidence. Nucleic Acids Res 46 (D1), D1062–D1067 (2018).

18. Polla, D.L. et al., Use of Targeted Exome Sequencing for Molecular Diagnosis of Skeletal Disorders. PLoS One 10 (9), e0138314 (2015).

19. Gibson, W.T. et al., Mutations in EZH2 cause Weaver syndrome. Am J Hum Genet 90 (1), 110–118 (2012).

20. Xie, H. et al., Polycomb repressive complex 2 regulates normal hematopoietic stem cell function in a developmental-stage-specific manner. Cell Stem Cell 14 (1), 68–80 (2014).

21. Velichutina, I. et al., EZH2-mediated epigenetic silencing in germinal center B cells contributes to proliferation and lymphomagenesis. Blood 116 (24), 5247–5255 (2010).

22. Kasinath, V. et al., Structures of human PRC2 with its cofactors AEBP2 and JARID2. Science 359 (6378), 940–944 (2018).

23. Poepsel, S., Kasinath, V., & Nogales, E., Cryo-EM structures of PRC2 simultaneously engaged with two functionally distinct nucleosomes. Nat Struct Mol Biol 25 (2), 154–162 (2018).

24. Sudlow, C. et al., UK biobank: an open access resource for identifying the causes of a wide range of complex diseases of middle and old age. PLoS Med 12 (3), e1001779 (2015).

25 All of Us Research Program, I., et al., The “All of Us” Research Program. N Engl J Med 381 (7), 668–676 (2019).

26. Shechter, D., Dormann, H.L., Allis, C.D., & Hake, S.B., Extraction, purification and analysis of histones. Nat Protoc 2 (6), 1445–1457 (2007).

27. Kasinath, V., Poepsel, S., & Nogales, E., Recent Structural Insights into Polycomb Repressive Complex 2 Regulation and Substrate Binding. Biochemistry 58 (5), 346–354 (2019).

28. Goering, R., Arora, A., Pockalny, M.C., & Taliaferro, J.M., RNA localization mechanisms transcend cell morphology. Elife 12 (2023).

29. Khandelia, P., Yap, K., & Makeyev, E.V., Streamlined platform for short hairpin RNA interference and transgenesis in cultured mammalian cells. Proc Natl Acad Sci U S A 108 (31), 12799–12804 (2011).

30. Kaya-Okur, H.S. et al., CUT&Tag for efficient epigenomic profiling of small samples and single cells. Nat Commun 10 (1), 1930 (2019).

31. Moses, J.T. et al., Neuro293: A REST-knockout HEK-293 cell line enables the expression of neuron-restricted genes for the high-throughput testing of human neurobiology and the biochemistry of neuronal proteins. Biol Methods Protoc 10 (1), bpaf036 (2025).

32. Hojfeldt, J.W. et al., Accurate H3K27 methylation can be established de novo by SUZ12-directed PRC2. Nat Struct Mol Biol 25 (3), 225–232 (2018).

33. Davidovich, C., Zheng, L., Goodrich, K.J., & Cech, T.R., Promiscuous RNA binding by Polycomb repressive complex 2. Nat Struct Mol Biol 20 (11), 1250–1257 (2013).

34. Lee, C.H. et al., Distinct Stimulatory Mechanisms Regulate the Catalytic Activity of Polycomb Repressive Complex 2. Mol Cell 70 (3), 435–448 e435 (2018).

35. Hojfeldt, J.W. et al., Non-core Subunits of the PRC2 Complex Are Collectively Required for Its Target-Site Specificity. Mol Cell 76 (3), 423–436 e423 (2019).

36. Glancy, E. et al., PRC2.1- and PRC2.2-specific accessory proteins drive recruitment of different forms of canonical PRC1. Mol Cell 83 (9), 1393–1411 e1397 (2023).

37. Sanulli, S. et al., Jarid2 Methylation via the PRC2 Complex Regulates H3K27me3 Deposition during Cell Differentiation. Mol Cell 57 (5), 769–783 (2015).

38. Nakagawa, R. et al., MicroRNA-155 controls affinity-based selection by protecting c-MYC+ B cells from apoptosis. J Clin Invest 126 (1), 377–388 (2016).

39. Ning, F. et al., Transcription factor Phf19 positively regulates germinal center reactions that underlies its role in rheumatoid arthritis. Am J Transl Res 10 (1), 200–211 (2018).

40 Bjeije, H., et al., Functional Comparison to Ezh2 Reveals PRC2-Independent Functions of Jarid2 in Hematopoietic Stem Cell Lineage Commitment. *bioRxiv* (2025).

41. Deevy, O. et al., Dominant-negative effects of Weaver syndrome-associated EZH2 variants. Genes Dev 39 (21-22), 1355–1376 (2025).

42. Bonilla, F.A. et al., Practice parameter for the diagnosis and management of primary immunodeficiency. J Allergy Clin Immunol 136 (5), 1186–1205 e1181-1178 (2015).

43. Bodenhofer, U., Bonatesta, E., Horejs-Kainrath, C., & Hochreiter, S., msa: an R package for multiple sequence alignment. Bioinformatics 31 (24), 3997–3999 (2015).

44. Thompson, J.D., Higgins, D.G., & Gibson, T.J., CLUSTAL W: improving the sensitivity of progressive multiple sequence alignment through sequence weighting, position-specific gap penalties and weight matrix choice. Nucleic Acids Res 22 (22), 4673–4680 (1994).

45. Riising, E.M. et al., Gene silencing triggers polycomb repressive complex 2 recruitment to CpG islands genome wide. Mol Cell 55 (3), 347–360 (2014).

46. Trouth, A. et al., The length of the G1 phase is an essential determinant of H3K27me3 landscapes across diverse cell types. PLoS Biol 23 (4), e3003119 (2025).

47. Langmead, B. & Salzberg, S.L., Fast gapped-read alignment with Bowtie 2. Nat Methods 9 (4), 357–359 (2012).

48. Danecek, P. et al., Twelve years of SAMtools and BCFtools. Gigascience 10 (2) (2021).

49. Ramirez, F. et al., deepTools2: a next generation web server for deep-sequencing data analysis. Nucleic Acids Res 44 (W1), W160–165 (2016).

50. Meers, M.P., Tenenbaum, D., & Henikoff, S., Peak calling by Sparse Enrichment Analysis for CUT&RUN chromatin profiling. Epigenetics Chromatin 12 (1), 42 (2019).

51. Ross-Innes, C.S. et al., Differential oestrogen receptor binding is associated with clinical outcome in breast cancer. Nature 481 (7381), 389–393 (2012).

52. Mudge, J.M. et al., GENCODE 2025: reference gene annotation for human and mouse. Nucleic Acids Res 53 (D1), D966–D975 (2025).

53. Yu, G., Wang, L.G., & He, Q.Y., ChIPseeker: an R/Bioconductor package for ChIP peak annotation, comparison and visualization. Bioinformatics 31 (14), 2382–2383 (2015).

54. Kramer, N.E. et al., Plotgardener: cultivating precise multi-panel figures in R. Bioinformatics 38 (7), 2042–2045 (2022).

55. Balas, M.M. et al., Establishing RNA-RNA interactions remodels lncRNA structure and promotes PRC2 activity. Sci Adv 7 (16) (2021).

