## Supplemental figures and Tables 1 and 2 for "Characterization of an immunodeficiency-associated EZH2 variant"

1. Supplemental Figures
    a. Supplemental Figure 1. EZH2 protein sequence alignments across several species.
    b. Supplemental Figure 2. Flow cytometry 1.
    c. Supplemental Figure 3. Flow cytometry 2.
    d. Supplemental Figure 4. Characterization of EZH2 expression.
    e. Supplemental Figure 5. CUT&Tag fragment size distribution.
    f. Supplemental Figure 6. Shared H3K27me3, EZH2, and FLAG CUT&Tag peaks in WT and L50S cell lines.
2. Supplemental Tables
    a. Supplemental Table 1. Cell lines used in the present study.
    b. Supplemental Table 2. Antibodies used in the present study

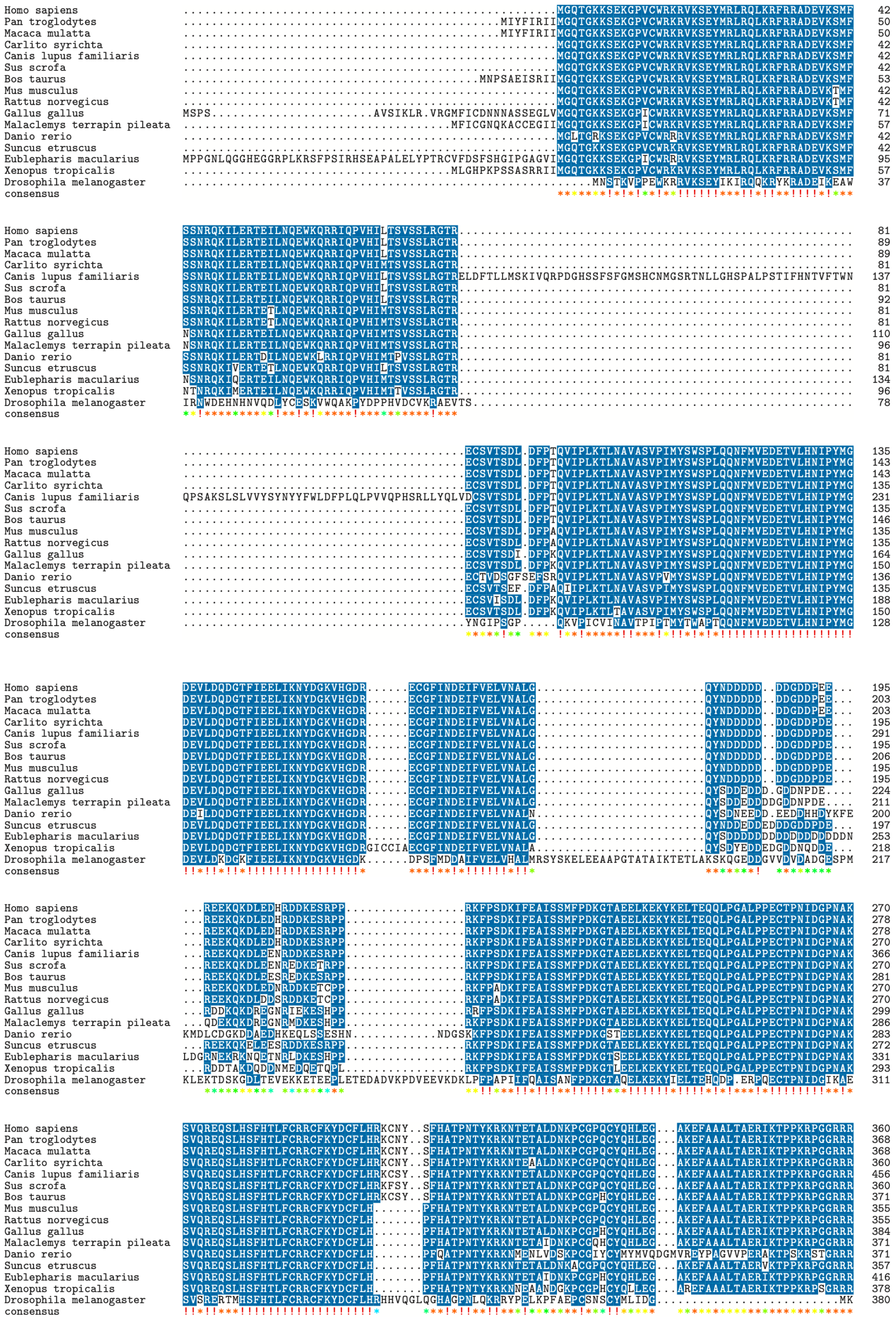

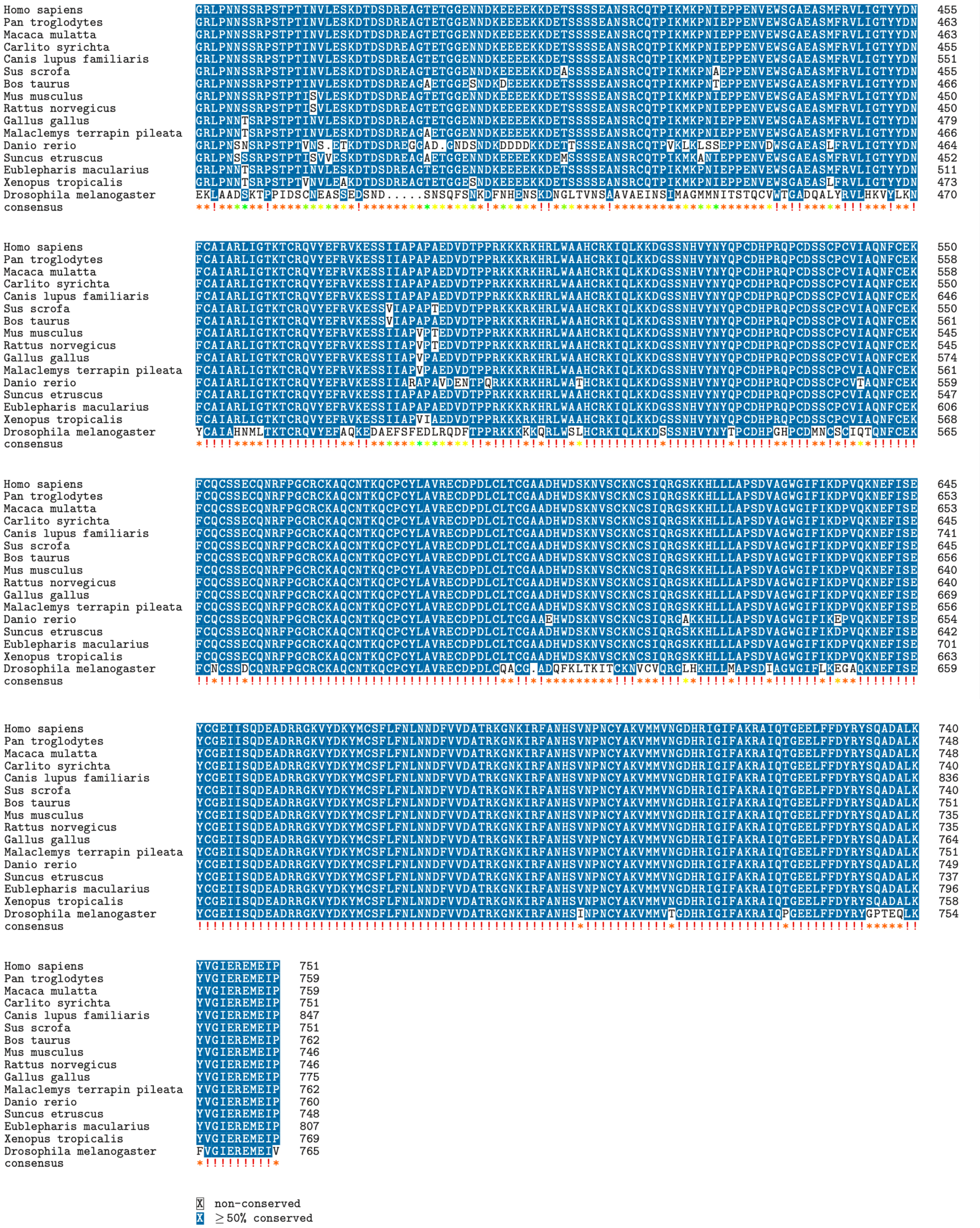


**Supplemental Figure 1. EZH2 protein alignment across several species.** Orthologous EZH2 protein sequences for all vertebrates were downloaded from *NCBI orthologs,* while *Drosophila melanogaster* ortholog sequence was downloaded from *GeneCards* orthologs database.

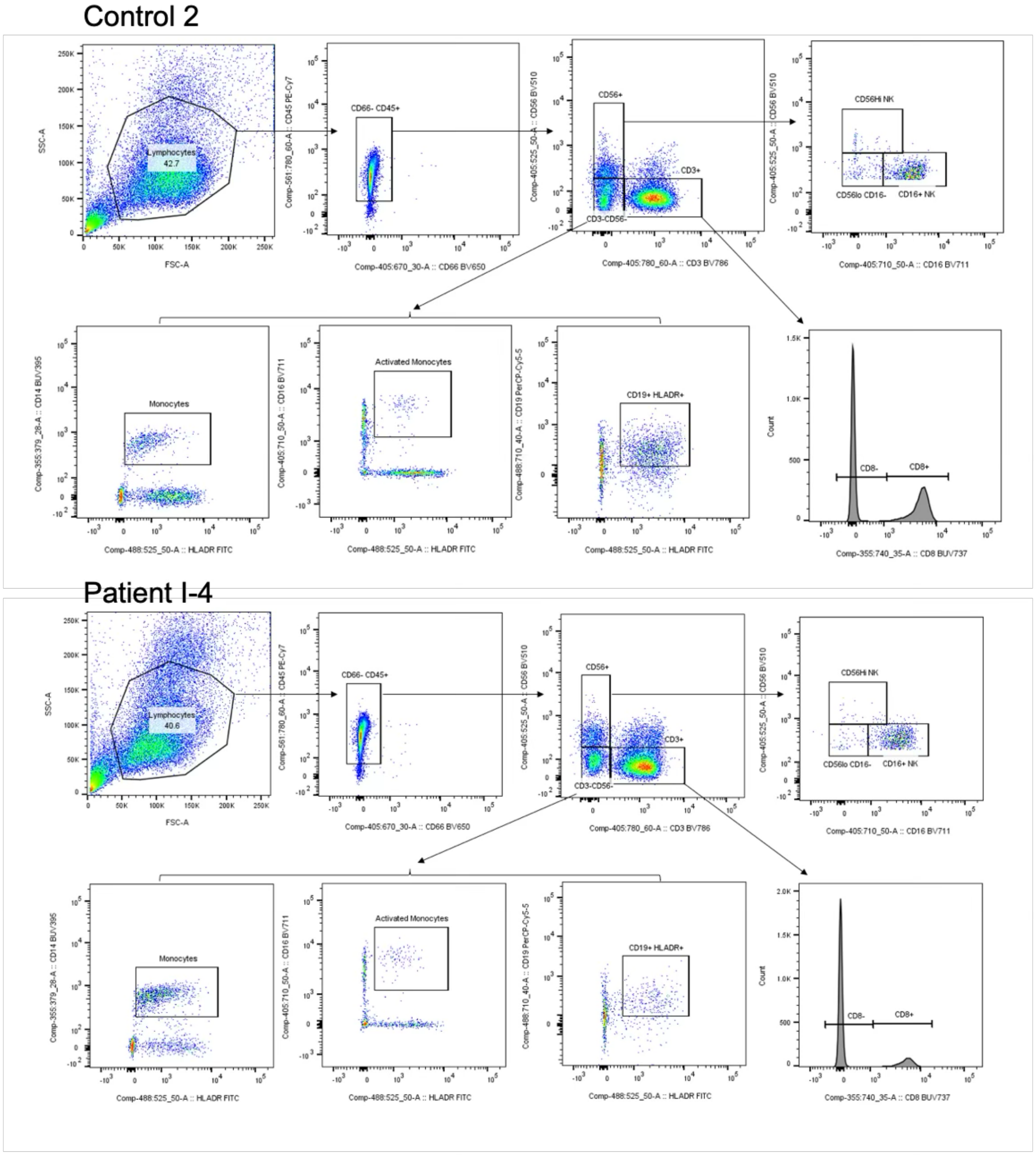


| | CD8 T cells | CD4 T cells | Monocytes | Activated Monoytes | B cells | CD56dim CD16+ NK cells | CD56high CD16- NK cells | CD56dim CD16dim NK cells |
|---|---|---|---|---|---|---|---|---|
| Control 2 | 31.1 | 42.2 | 3 | 0.46 | 6.54 | 5.15 | 0.25 | 0.67 |
| Patient I-4 | 13.2 | 59.4 | 7.18 | 0.52 | 1.45 | 5.47 | 0.19 | 0.34 |

*Frequencies of PBMC subsets shown as a frequency of CD45+ CD66- leukocytes

**Supplemental Figure 2. Flow cytometry 1.** Gating strategies shown for “Control 2” and Patient “I-4” leukocyte samples with percentages of each cell type shown below.

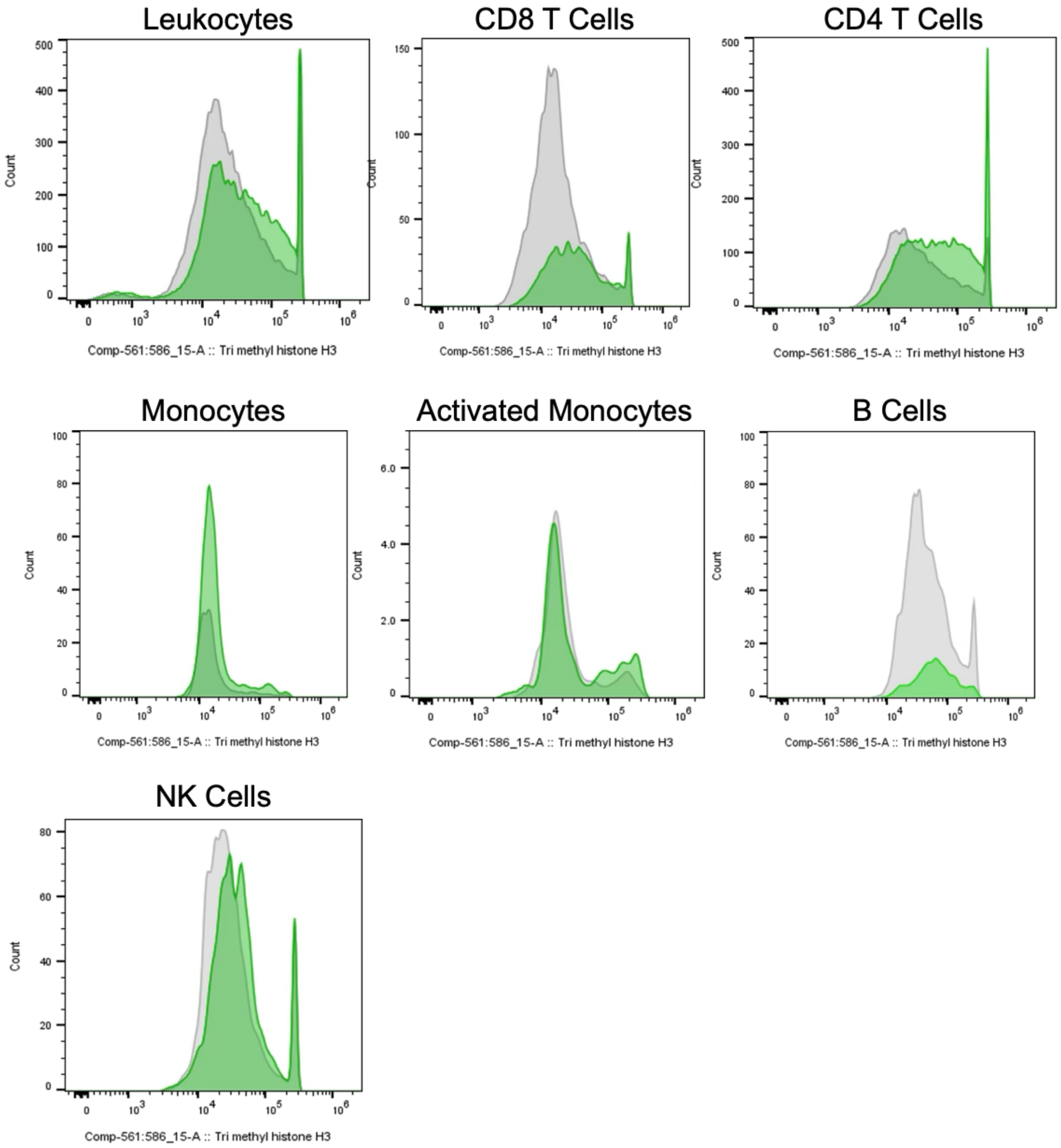


| | Leukocytes | CD8 T Cells | CD4 T Cells | Monocytes | Activated Monocytes | B Cells | NK Cells |
|---|---|---|---|---|---|---|---|
| Control | 46749 | 32544 | 49438 | 24381 | 38167 | 62656 | 47662 |
| Patient B014 | 73572 | 62565 | 85366 | 26345 | 56556 | 77313 | 56967 |
| FMO | 1810 | 1448 | 2118 | 1445 | 2402 | 1860 | 1287 |

*Mean fluorescence intensities of trimethyl histone H3 antibody staining shown for the indicated PBMC subsets. FMO = fluorescence-minus-one control

**Supplemental Figure 3. Flow cytometry 2.** Intensity values for histone H3 lysine 27 trimethylation staining .

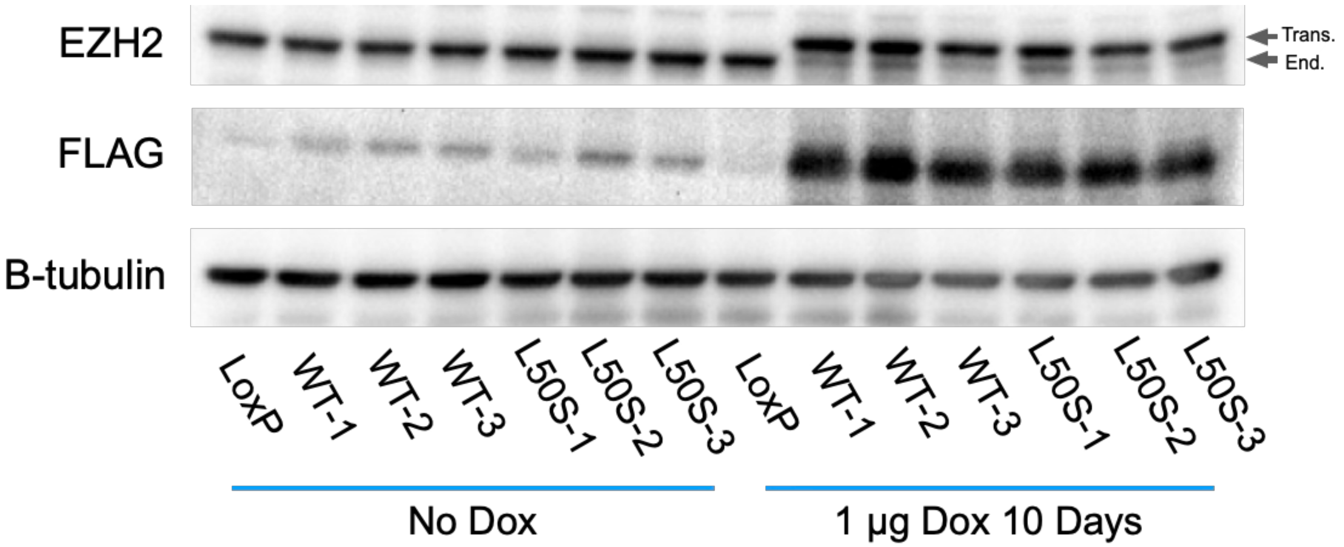


**Supplemental Figure 4. Characterization of EZH2 transgene expression.** Doxycycline-inducible EZH2 constructs were introduced into HEK-293T cell lines engineered with a LoxP-containing safe harbor locus. Top, endogenous EZH2 expression under non-inducible conditions (No Dox) and doxycycline inducible (Dox) conditions. Middle, FLAG-tagged EZH2 expression under non-inducible and inducible conditions. Bottom, B-tubulin as loading control. Endogenous EZH2 (End.), transgene EZH2 (Trans).

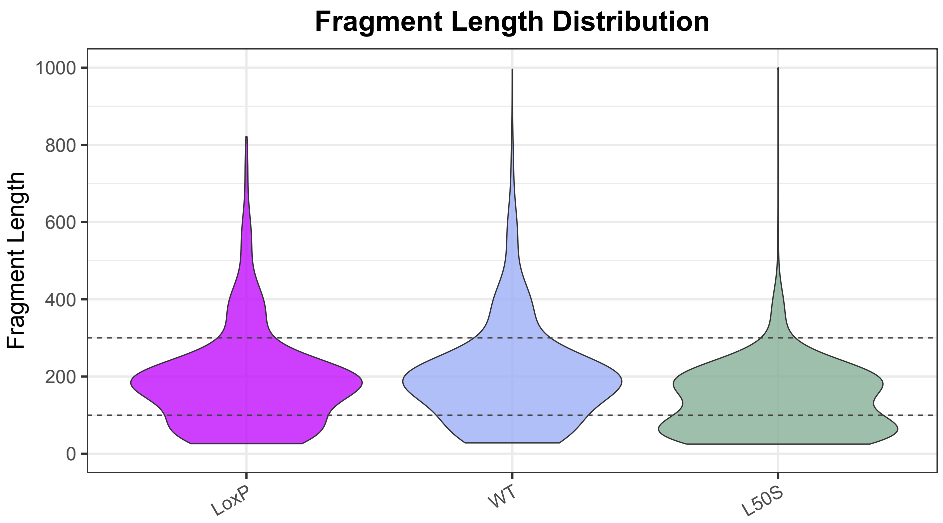


**Supplemental Figure 5. CUT&Tag fragment size distribution.** Fragment size distribution before alignment sieve between 100 bp and 300bp (indicated by dashed lines).

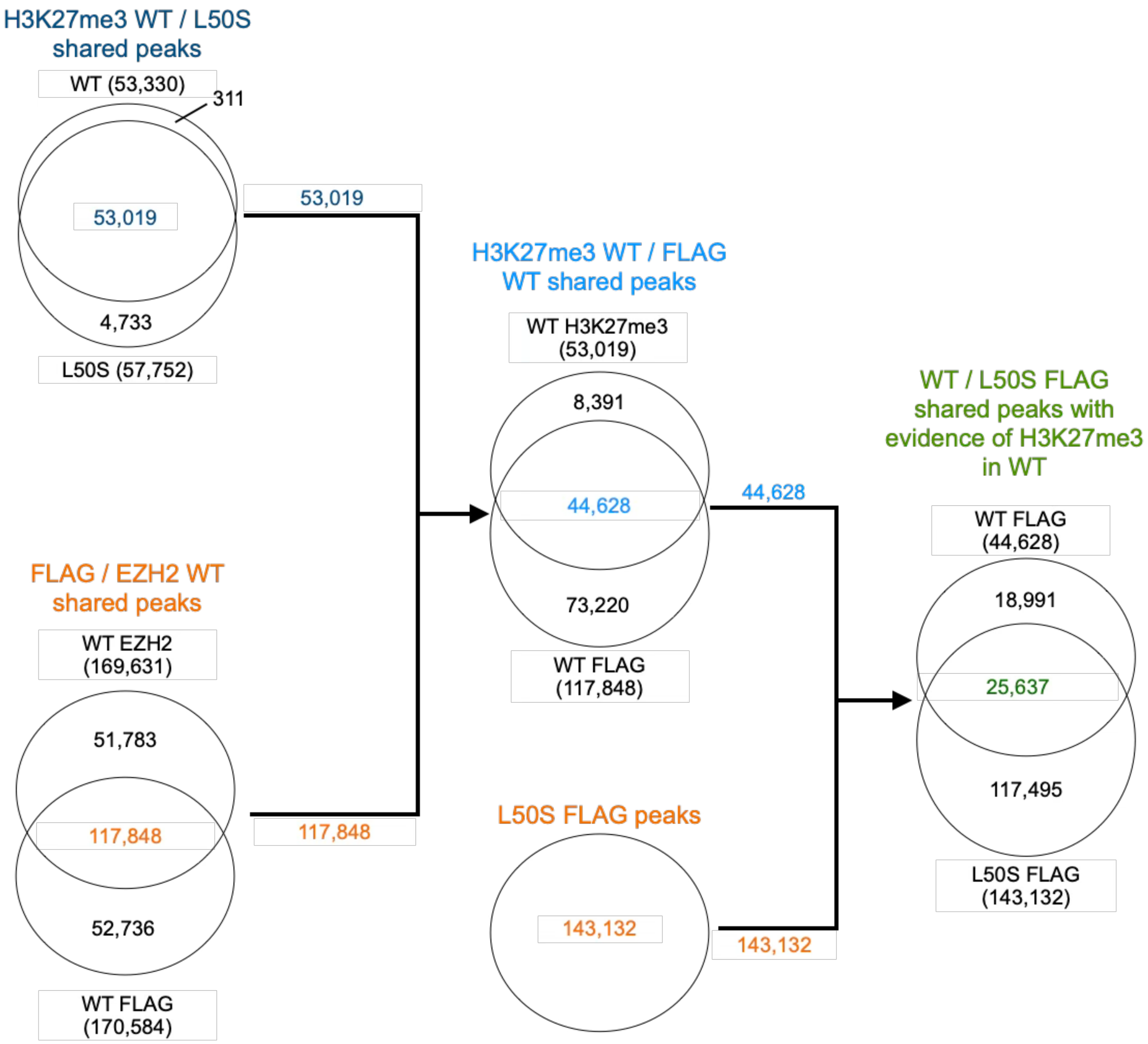


**Supplemental Figure 6. Shared H3K27me3, EZH2, and FLAG CUT&Tag peaks in WT and L50S cell lines.** Venn diagrams showing overlapping peaks between merged biological replicates of WT and L50S cell lines in H3K27me3, EZH2 and L50S CUT&Tag experiments.

| Cell Line | Parental Cell Line | Relevant Genotype | Source | Citation |
|---|---|---|---|---|
| LoxP | HEK293T | Parental EZH2 only endogenous | Matthew Taliaferro Laboratory | Khandelia et al. (2011) PMID:21768390 |
| WT-1 | LoxP | Parental EZH2 endogenous / Doxycycline inducible WT EZH2 transgene | Generated in this study | Present study |
| WT-2 | LoxP | Parental EZH2 endogenous / Doxycycline inducible WT EZH2 transgene | Generated in this study | Present study |
| WT-3 | LoxP | Parental EZH2 endogenous / Doxycycline inducible WT EZH2 transgene | Generated in this study | Present study |
| L50S-1 | LoxP | Parental EZH2 endogenous / Doxycycline inducible L50S EZH2 transgene | Generated in this study | Present study |
| L50S-2 | LoxP | Parental EZH2 endogenous / Doxycycline inducible L50S EZH2 transgene | Generated in this study | Present study |
| L50S-3 | LoxP | Parental EZH2 endogenous / Doxycycline inducible L50S EZH2 transgene | Generated in this study | Present study |
| GM12878 | N/A | Parental EZH2 only endogenous | Srinivas Ramachandran Laboratory | RRID:CVCL_7526 |

**Supplemental Table.1 Cell lines used in the present study.**

Supplemental Materials

| | Antibody | Fluorochrome | Clone | Company | Catalog # |
|---|---|---|---|---|---|
| **Flow Cytometry** | CD3 | BV786 | UCHT1 | BD Bioscience | 565491 |
| | CD4 | PE-Dazzle | SK3 | Biolegend | 344640 |
| | CD8 | BUV737 | SK1 | BD Bioscience | 564629 |
| | CD66 | BV650 | B1.1/CD66 | BD Bioscience | 740607 |
| | CD19 | PerCP-Cy5.5 | HIB19 | Biolegend | 302229 |
| | CD56 | BV510 | HCD56 | Biolegend | 318339 |
| | CD11c | APC-Cy7 | Bu15 | Biolegend | 337218 |
| | CD16 | BV711 | 3G8 | Biolegend | 302044 |
| | CD11b | BV421 | ICRF44 | ThermoFisher | 62011841 |
| | CD45 | PE-Cy7 | HI30 | Biolegend | 304015 |
| | CD15 | PE-Cy5 | W6D3 | Biolegend | 323033 |
| | HLADR | FITC | L243 | ThermoFisher | 11-9952-41 |
| | CD14 | BUV395 | M5E2 | BD Bioscience | 740286 |
| | H3K27me3 | PE | C36B11 | Cell Signaling Technologies | 40724S |
| **Immunoblotting** | H3K27me3 | N/A | C36B11 | Cell Signaling Technologies | 9733 |
| | H3K27me2 | N/A | D18C8 | Cell Signaling Technologies | 9728 |
| | H3K27me1 | N/A | MABI0321 | Active Motif | 61016 |
| | EZH2 | N/A | D2C9 | Cell Signaling Technologies | 5246 |
| | FLAG | N/A | M2 | Sigma-Aldrich | F1804 |
| | SUZ12 | N/A | D39F6 | Cell Signaling Technologies | 3737 |
| | EED | N/A | E4L6E | Cell Signaling Technologies | 85322 |
| | B Tubulin | N/A | BT7R | UBPBio | Y1061 |

| | | | | | |
|---|---|---|---|---|---|
| | Goat Anti-Mouse IgG – HRP Conjugated | N/A | - | BIO-RAD | 1706516 |
| | Goat Anti-Rabbit IgG – HRP Conjugated | N/A | - | BIO-RAD | 1705046 |
| **CUT&Tag** | H3K27me3 | N/A | C36B11 | Cell Signaling Technologies | 9733 |
| | EZH2 | N/A | D2C9 | Cell Signaling Technologies | 5246 |
| | FLAG | N/A | M2 | Sigma-Aldrich | F1804 |
| | IgG | N/A | - | EpiCypher | 13-0042 |
| | Anti-rabbit | N/A | - | EpiCypher | 13-0047 |
| | Anti-mouse | N/A | - | EpiCypher | 13-0048 |

**Supplemental Table 2. Antibodies used in the present study.** N/A not applicable. – No information available.
